# A large-scale neural network training framework for generalized estimation of single-trial population dynamics

**DOI:** 10.1101/2021.01.13.426570

**Authors:** Mohammad Reza Keshtkaran, Andrew R. Sedler, Raeed H. Chowdhury, Raghav Tandon, Diya Basrai, Sarah L. Nguyen, Hansem Sohn, Mehrdad Jazayeri, Lee E. Miller, Chethan Pandarinath

## Abstract

Recent technical advances have enabled recording of increasingly large populations of neural activity, even during natural, unstructured behavior. Deep sequential autoencoders are the current state-of-the-art for uncovering dynamics from these datasets. However, these highly complex models include many non-trainable hyperparameters (HPs) that are typically hand tuned with reference to supervisory information (e.g., behavioral data). This process is cumbersome and time consuming and biases model selection toward models with good representations of individual supervisory variables. Additionally, it cannot be applied to cognitive areas or unstructured tasks for which supervisory information is unavailable. Here we demonstrate AutoLFADS, an automated model-tuning framework that can characterize dynamics using only neural data, without the need for supervisory information. This enables inference of dynamics out-of-the-box in diverse brain areas and behaviors, which we demonstrate on several datasets: motor cortex during free-paced reaching, somatosensory cortex during reaching with perturbations, and dorsomedial frontal cortex during cognitive timing tasks. We also provide a cloud software package and comprehensive tutorials that enable new users to apply the method without dedicated computing resources.

## Introduction

Ongoing advances in neural interfacing technologies are enabling simultaneous monitoring of the activity of large neural populations across a wide array of brain areas and behaviors (1–5). Such technologies may fundamentally change the questions we can address about computations within a neural population, allowing neuroscientists to shift focus from understanding how individual neurons’ activity relates to externally-measurable or controllable parameters, toward understanding how neurons within a network coordinate their activity to perform computations underlying those behaviors. A natural framework for interpreting these complex, high-dimensional datasets is through modeling the neural population’s dynamics (6–8). The dynamical systems framework centers on uncovering coordinated patterns of activation across a neural population and characterizing how these patterns change over time. Knowledge of these hidden dynamics has provided new insights into how neural populations implement the computations necessary for motor, sensory, and cognitive processes (9–15).

A focus on population dynamics could also facilitate a shift away from reliance on stereotyped behaviors and trial-averaged neural responses. Standard approaches must typically average activity across trials, sacrificing single trial interpretability for robustness against what is perceived as noise in single trials. However, as articulated by Cunningham and Yu (16): “If the neural activity is not a direct function of externally measurable or controllable variables (for example, if activity is more a reflection of internal processing than stimulus drive or measurable behavior), the time course of neural responses may differ substantially on nominally identical trials.” This may be especially true of non-primary cortical areas, and cognitively demanding tasks that involve decision-making, allocation of attention, or varying levels of motivation.

Several approaches have been developed to infer latent dynamical structure from neural population activity on individual trials, including a growing number that leverage artificial neural networks (17–22). One such method, latent factor analysis via dynamical systems (LFADS) (20,22) achieved precise inference of motor cortical firing rates on single trials of stereotyped behaviors, enabling accurate prediction of subjects’ behaviors on a moment-by-moment, millisecond timescale (20). Further, in tasks with unpredictable events, a modified network architecture enabled inference of dynamical perturbations that corresponded to how subjects ultimately responded to the unpredictable events.

Though highly effective, artificial neural networks typically have many thousands of parameters, and potentially dozens of non-trainable hyperparameters (HPs) that control architecture, regularization, and parameter optimization. Until recently, HP optimization consisted of an iterative manual process, a random search, or some combination of the two. In the past several years, a host of more advanced approaches promises to eliminate the tedious work and domain knowledge required for manual tuning while performing better and more efficiently than random search (23–25). The form and variety of neuroscientific datasets present unique challenges that make HP optimization a particularly impactful problem (26). Thus, bringing efficient HP search algorithms to neuroscience could allow more effective experimentation with models based on artificial neural networks, like LFADS.

Here we present AutoLFADS, a framework for large-scale, automated model tuning that enables accurate single-trial inference of neural population dynamics across a range of brain areas and behaviors. We evaluate AutoLFADS using data from three cortical regions: primary motor and dorsal premotor cortex (M1/PMd), somatosensory cortex area 2, and dorsomedial frontal cortex (DMFC). The tasks span a mix of functions where population activity can be well-modeled by autonomous dynamics (e.g., pre-planned reaching movements, estimation of elapsed time) and those for which population activity is responsive to external inputs (e.g., mechanical perturbations, unexpected appearance of reaching targets, variable timing cues).

Using this broad range of datasets, we show that AutoLFADS achieves high-time resolution, single-trial inference of neural population dynamics, surpassing LFADS in all scenarios tested. Remarkably, AutoLFADS does this in a completely unsupervised manner that does not depend on the knowledge of the tasks, subjects’ behaviors, or brain areas. In all applications, the method is applied “out of the box” without careful adjustment for each dataset. These capabilities greatly extend the range of neuroscientific applications for which accurate inference of single-trial population dynamics should be achievable, and substantially lower the barrier to entry for applying these methods. Finally, we present a cloud software package and comprehensive tutorials to enable new users without machine learning expertise or dedicated computing resources to apply AutoLFADS successfully.

## Results

### LFADS training

The LFADS architecture (**Fig. 1a**) has been detailed previously (20,22,26), and we give a brief overview in *Methods*. LFADS is a variant of sequential variational autoencoder (VAE) (22,27,28) that approximates the latent dynamical system underlying the observed population using a recurrent neural network (RNN; the “Generator”) and a sequence of inferred inputs. The model receives sequences of binned spike counts as input and produces a Poisson firing rate for each neuron and time step. When training the model, the objective is to maximize a lower bound on the marginal likelihood of the observed spiking activity given the inferred rates (see *Methods* for details).

**Figure 1.**
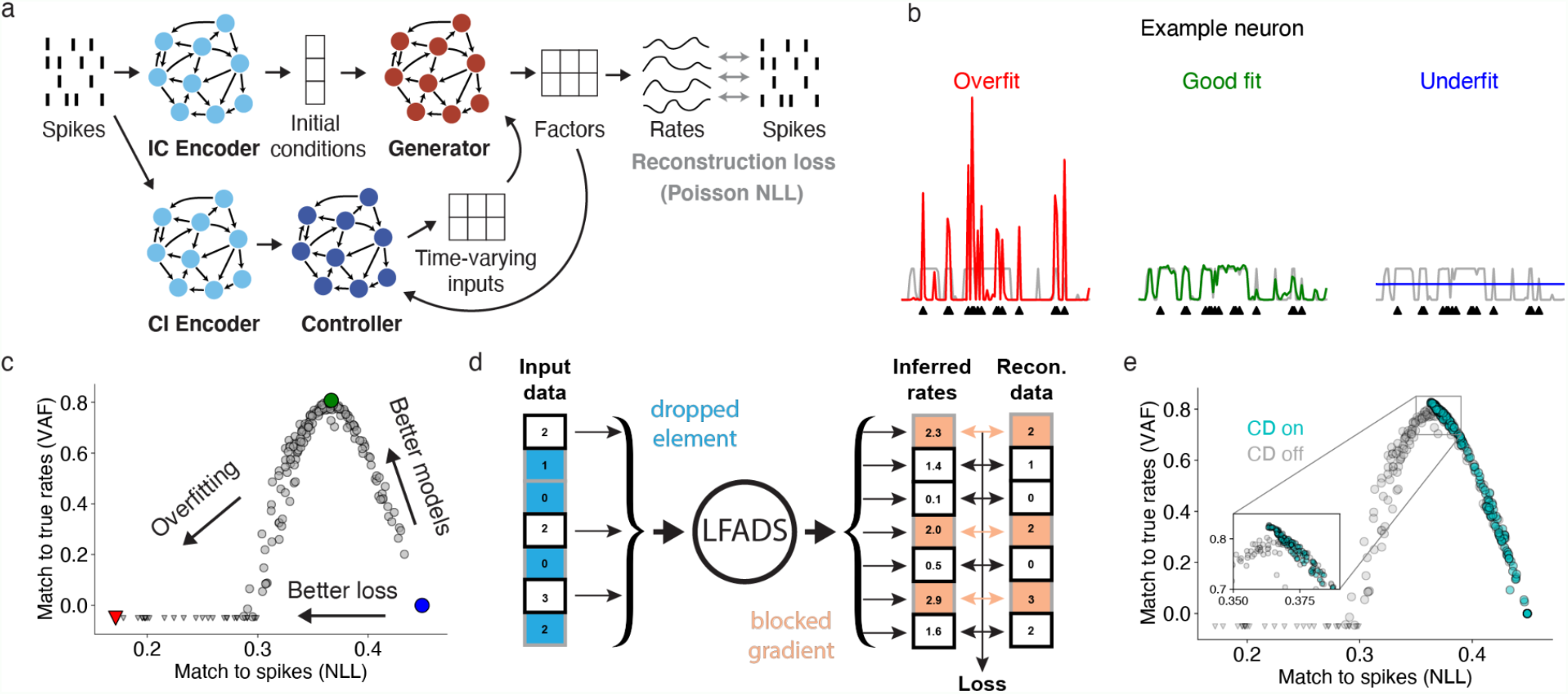
AutoLFADS combines a novel neural network regularization method with a large-scale framework for automated hyperparameter optimization. (a) Schematic of the LFADS architecture, showing how the generative model infers the firing rates that underlie the observed spikes. (b) Examples of LFADS-inferred rates (colored), the corresponding synthetic input data (spikes, shown as black triangles), and the data-generating distribution (ground truth rates, shown as gray traces) for three fitting modes. (c) Performance of 200 LFADS models with random HPs in matching the spikes and the known rates of a synthetic dataset, measured by validation negative log-likelihood (NLL) and variance accounted for (VAF) respectively. Gray points indicate random search models and colored points indicate the models that produced the rates in the previous panel. Triangles indicate models with negative VAF; actual values were ignored for better visualization. All metrics were computed on validation data. (d) Schematic of CD. Random elements of the input data tensor are zeroed and the rest are scaled up, as in the standard dropout layer. Loss gradients are blocked for these elements to prevent overfitting to spikes. (e) Same as in (c), but for models trained with CD.

It is imperative to regularize the model properly in order to extract denoised firing rates (**Fig. 1b**) (26). This can be achieved through HP optimization. The two main classes of LFADS HPs are those that set the network architecture (e.g., number of units in each RNN, dimensionality of initial conditions, inputs, and factors), and those that control regularization and training (e.g., L2 penalties, scaling factors for KL penalties, dropout probability, and learning rate; described in *Methods*). The optimal values of these HPs could depend on various factors such as dataset size, dynamical structure underlying the activity of the brain region being modeled, and the behavioral task.

A critical challenge for autoencoders is that automatic HP searches face a type of failure mode that is particularly hard to address (26). Given enough capacity, the model can find a trivial solution where it simply passes individual spikes from the input to the output firing rates, akin to an identity transformation of the input, without modeling any meaningful structure underlying the data (**Fig. 1b**, left). Importantly, this failure mode cannot be detected via the standard strategy of evaluating validation loss on held-out trials, because the identity-like transformation performs similarly on training and validation data. We term this failure mode “identity overfitting” and distinguish it from the standard notion of overfitting to the training data. We performed a 200-model random search over a space of KL, L2, and dropout regularization HPs that was empirically determined to yield both underfit and identity-overfit models on a synthetic dataset with known firing rates (see *Methods* for a description of the dataset). Models that appear to have the best likelihoods actually exhibit poor inference of underlying firing rates, a hallmark of identity overfitting (**Fig. 1c**). This phenomenon is also consistently observed on real data throughout this paper: better validation loss did not indicate better performance for any of our decoding or PSTH-based metrics.

The lack of a reliable validation metric has prevented automated HP searches because it is unclear how one should select between models when underlying firing rates are unavailable or non-existent. While comparing likelihoods for data seen and not seen by the encoders (e.g., via co-smoothing (29) or sample validation (26)) can give some indication of the degree of identity overfitting, these metrics alone do not provide sufficient information for model selection. To address this issue, we developed a novel regularization technique called coordinated dropout (CD; (26)). CD operates based on the reasonable assumption that the observed high-dimensional neural activity originates from a latent, lower dimensional subspace (30,31). CD forces the network to model only structure that is shared across neurons by regularizing the flow of information through the network during training. At each training step, a random mask is applied to drop data elements (spike count bins, across time and neurons) at the network’s input, while scaling up spike counts so the expected sum of the input remains the same (**Fig. 1d**). The complement of this mask is applied at the network’s output to block the gradient flow from the samples that are present at the input. Thus, the network cannot learn to use a neuron’s activity at a given time step to predict its own firing rate at that time step. This strategy effectively prevents the network from learning a trivial identity transformation because individual data samples are never used for self reconstruction. Note that CD is a regularization strategy and thus is only applicable during training to regularize the optimization.

We repeated the previous test on synthetic data by training 200 LFADS models from the same HP search space with CD, and found that they no longer overfit spikes (**Fig. 1e**). CD, in a completely unsupervised way, restored the correspondence between model quality assessed from matching spikes (validation likelihood) and matching rates, allowing the former to be used as a surrogate when the latter is not available. By preventing identity overfitting, CD renders the validation likelihood a reliable metric for model selection. Summary metrics for these and all subsequent analyses are aggregated in **Supp. Table 1**.

The premise of this paper is that this reliable validation metric should enable large-scale HP searches and unsupervised, fully-automated selection of high-performing neuroscientific models despite having no access to ground truth firing rates. To test this, we needed an efficient HP search strategy. We chose a recent method based on parallel search called Population Based Training (PBT; **Supp. Fig. 1**) (25,32). PBT distributes training across dozens of workers simultaneously, and uses evolutionary algorithms to tune HPs over many generations. Because PBT distributes model training over many workers, it matches the scalability of parallel search methods such as random or grid search, while achieving higher performance with the same amount of computational resources (**Supp. Fig. 1**) (25,32). PBT also enables dynamic HP adjustment throughout the course of training and allows exploration outside the search initialization ranges (**Supp. Fig. 1**).

These two key modifications - a novel regularization strategy (CD) that results in a reliable validation metric, and an efficient approach to HP optimization (PBT) - yield a large-scale, automated framework for model tuning, which we refer to as AutoLFADS. In the following sections, we characterize the performance of AutoLFADS on a diverse array of datasets, representing a variety of task structures and brain areas.

### AutoLFADS outperforms original LFADS when applied on benchmark data from M1/PMd

We first evaluated AutoLFADS on the delayed-reaching dataset used to develop and assess the original LFADS method (20). During this maze reaching task (see *Methods*), a monkey made a variety of straight and curved reaches after a delay period following target presentation (**Fig. 2a**), with multiple highly stereotyped trials per condition. Neural activity during preparation and reaching was recorded from M1 and PMd. Previous analyses of the delayed reaching paradigm demonstrated that activity during the movement period is well modeled by autonomous dynamics (10,20); i.e., the temporal evolution of the neural population’s activity is predictable based on the state it reaches during the delay period. Therefore, previous work modeled these data with a simplified LFADS configuration which could only approximate autonomous dynamics (20). In this paper we do not constrain the network architecture to only model autonomous dynamics for any applications tested, to determine whether AutoLFADS can automatically adjust the degree to which autonomous dynamics and inputs are needed to model the data.

**Figure 2.**
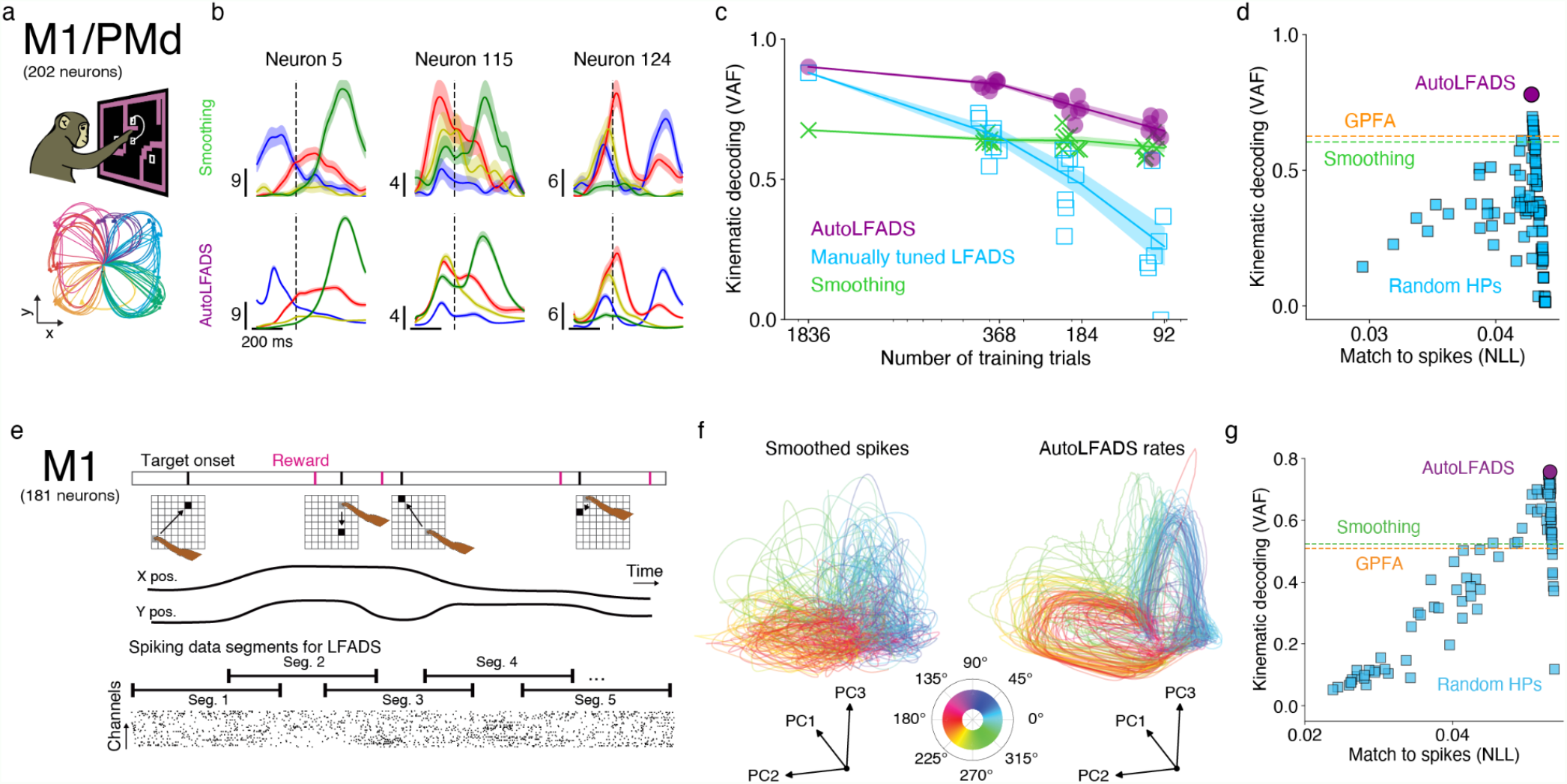
Application of AutoLFADS to data from motor cortex. Results are shown for the maze task (a-d) and the random target task (e-g). (a) Schematic of the maze task and representative reach trajectories across 108 conditions, colored by target location. (b) Example PSTHs, aligned to movement onset (dashed line). Colors indicate reach conditions, shaded regions denote standard error, and vertical scale bars denote rates (spikes/s). (c) Hand velocity decoding performance (VAF, mean of x- and y-directions) from firing rates as a function of training dataset size. Lines and shading denote mean +/- standard error across 7 models trained on randomly-drawn trial subsets. (d) Hand velocity decoding performance from a 184-trial training set in comparison to smoothing, GPFA, and LFADS random search (100 models; no CD) baselines. (e) Schematic of the random target task (top), showing a lack of stereotyped trial structure and delay periods. Continuous neural spiking data was divided into overlapping segments for modeling (bottom). After modeling, the inferred rates were merged using a weighted average of data at overlapping timepoints. (f) Subspaces of neural activity, separately extracted using PCA and colored by angle to the target. (g) Same as in (d), but for the random target dataset.

Consistent with previous applications of LFADS on this dataset (20,26), the firing rates inferred by AutoLFADS for 2 ms bins exhibited clear and consistent structure on individual trials (**Supp. Fig. 2**). We also verified that these firing rates captured features of the neural responses revealed by averaging smoothed spikes across trials, a common method of de-noising neural activity (**Fig. 2b**).

A generalizable method should be able to perform well across the broad range of dataset sizes typical of neuroscience experiments. To test this, we compared AutoLFADS and manually-tuned LFADS models that were trained using either the full dataset (2296 trials), or randomly sampled subsets containing 5, 10, and 20% of the trials. We quantified model performance by decoding the monkey’s hand velocity from the inferred rates using optimal linear estimation (**Fig. 2c**).

At the largest dataset size, decoding performance for AutoLFADS and manually-tuned LFADS was comparable. This result fits with standard intuition that performance is less sensitive to HPs when sufficient data are available. However, for all three reduced dataset sizes, AutoLFADS outperformed the manually-tuned model (p<0.05 for all three sizes, paired, one-tailed Student’s t-test).

While this result is promising, the difference in robustness to dataset size between AutoLFADS and LFADS could have resulted from a particularly poor selection of HPs during manual tuning. To control for this possibility, we trained 100 additional LFADS models with randomly-selected HPs on one of the smaller data subsets (184 trials). We evaluated the models’ performance using hand velocity decoding (**Fig. 2d**) as described above and reproduction of empirical trial-averaged firing rates (PSTHs; **Supp. Fig. 2**). As before, AutoLFADS maintains two advantages over LFADS with random search: completely unsupervised model selection and higher peak performance by both supervision metrics.

### AutoLFADS uncovers population dynamics without structured trials

To-date, most efforts to tie dynamics to neural computations have used experiments where subjects perform constrained tasks with repeated, highly structured trials. For example, motor cortical dynamics are often framed as a computational engine to link the processes of motor preparation and execution (6–8). To interrogate these dynamics, most studies use a delayed-reaching paradigm that creates explicit pre-movement and movement periods. However, constrained behaviors may have multiple drawbacks in studying dynamics. First, it is unclear whether such artificial paradigms are good proxies for everyday behaviors. Second, highly constrained, repeated behaviors might impose artificial limits on the properties of the uncovered dynamics, such as the measured dimensionality of the neural population activity (33). Even outside of movement neuroscience, the requirement that we conduct many repetitions of constrained tasks significantly hinders our ability to study a rich sample of the dynamics of a given neural population. Accurate inference of neural dynamics without these constraints could facilitate dynamics-based analyses of richer datasets that are more reflective of the brain’s natural behavior.

We applied AutoLFADS to neural activity from a monkey performing a continuous, self-paced random target reaching task (**Fig. 2e**, top) (34), in which each movement started and ended at a random position, and movements were highly variable in duration. Analysis of data without consistent temporal structure repeated across trials is challenging, as trial-averaging is not feasible. Further, the strong simplifying assumptions that are typically used for single-trial analyses - for example, that the arm is in a consistent starting point, and that data windows are aligned to behavioral events such as target or movement onset (17,19,26,35–39) - are not applicable or severely limiting when analyzing less-structured tasks.

Since typical fixed-length, trial-aligned data segments were not an option for this dataset, we chopped an approximately 9-minute window of continuous neural data into 600 ms segments with 200 ms of overlap (**Fig. 2e**, bottom) without regard to trial boundaries. After modeling with AutoLFADS, we merged inferred firing rates from individual segments, which yielded inferred rates for the original continuous window. We then analyzed the inferred rates by aligning the data to movement onset for each trial (see *Methods*). Even though the dataset was modeled without the use of trial information, inferred firing rates during the reconstructed trials exhibited consistent progression in an underlying state space, with clear structure that corresponded with the monkey’s reach direction on each trial (**Fig. 2f**, right). Again, we highlight the advantages of AutoLFADS in unsupervised model selection and higher peak performance (**Fig. 2g**). Further, we show that AutoLFADS learns to infer rates that clearly reveal velocity, position, and target subspaces without any information about task structure (**Supp. Note 1 & Supp. Fig. 3**).

### AutoLFADS accurately captures single-trial population dynamics in somatosensory cortex

Area 2 provides a valuable test case for generalization of AutoLFADS to input-driven brain areas. As a sensory area, area 2 receives strong afferent input from cutaneous receptors and muscles and is robustly driven by mechanical perturbations to the arm (35,40,41). Functionally, area 2 is thought to serve a role in mediating reach-related proprioception (35–37,41), was recently shown to contain information about whole-arm kinematics (35), and may also receive efferent input from motor areas (35,38,39,41). This macaque area 2 dataset consisted of recordings taken during active and passive arm movements in one of eight directions (**Fig. 3a**; see *Methods* for details).

**Figure 3.**
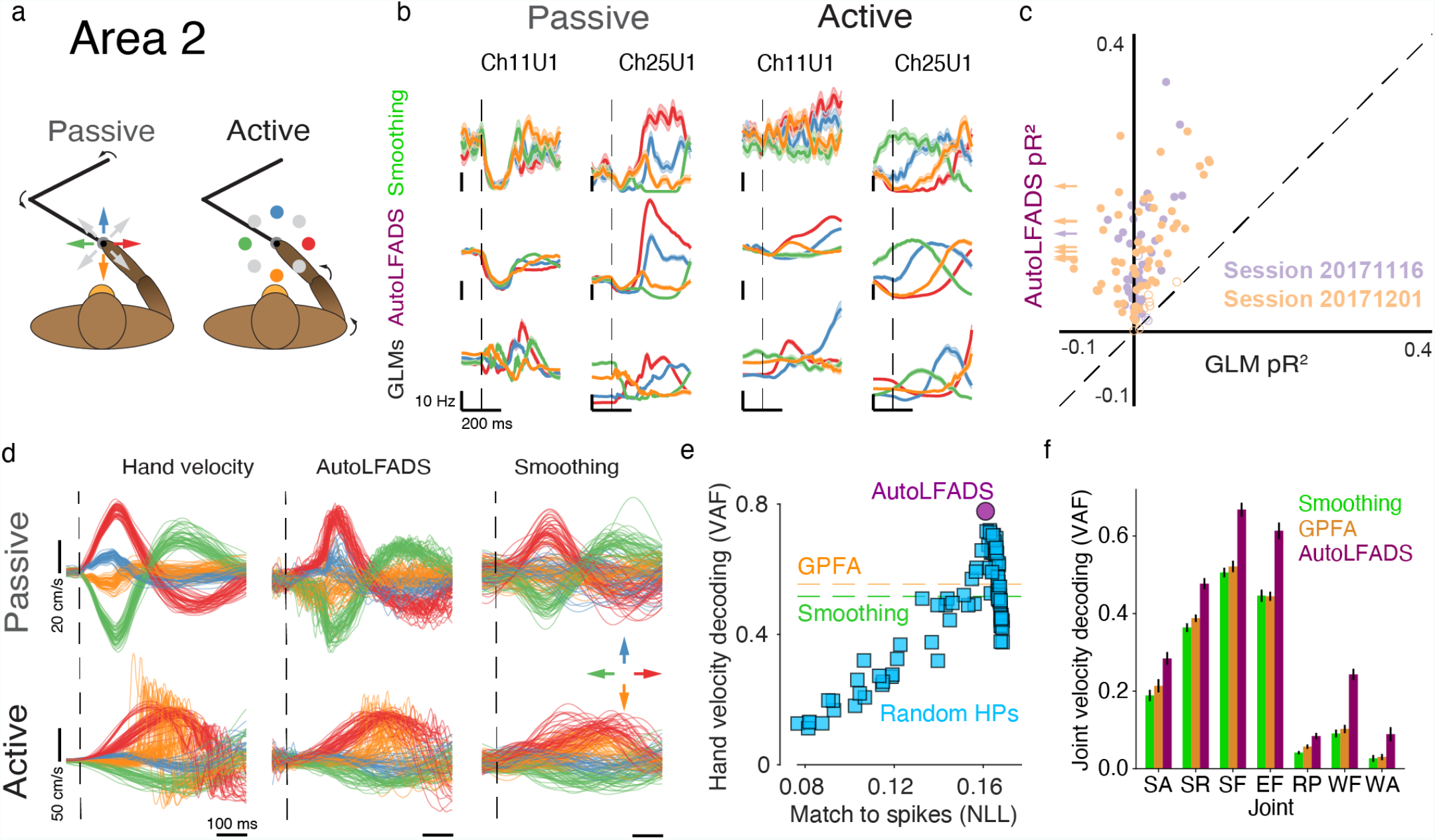
Application of AutoLFADS to data from somatosensory cortex area 2. (a) Schematic of the center-out, bump task showing passive and active conditions. (b) PSTHs produced by smoothing spikes (top), AutoLFADS (middle), or GLM predictions (bottom) for 3 example neurons. (c) Comparison of spike count predictive performance for AutoLFADS and GLMs. Filled circles correspond to neurons for which AutoLFADS pR^2^ was significantly higher than GLM pR^2^, and open circles correspond to neurons for which there was no significant difference. Arrows (left) indicate neurons for which GLM pR^2^ was outside of the plot bounds. (d) Subspace representations of hand x-velocity during active and passive movements extracted from smoothed spikes and rates inferred by AutoLFADS. (e) Comparison of AutoLFADS vs. random search (no CD) in decoding hand velocity during active trials. (f) Joint angular velocity decoding performance from firing rates inferred using smoothing, Gaussian process factor analysis (GPFA), and AutoLFADS. Error bars denote standard error of the mean. Joint abbreviations: shoulder adduction (SA), shoulder rotation (SR), shoulder flexion (SF), elbow flexion (EF), wrist radial pronation (RP), wrist flexion (WF), and wrist adduction (WA).

The single-trial rates inferred by AutoLFADS for passive trials exhibited clear and structured responses to the unpredictable perturbations (**Supp. Fig. 2**), highlighting the model’s ability to approximate input-driven dynamics.The inferred rates also had a close correspondence to the empirical PSTHs during active and passive trials (**Fig. 3b**), outperforming all LFADS models trained with random HPs (**Supp. Fig. 2**; passive only). Additionally, the inferred inputs show structure consistent with the supervised notions of trials, directions, and perturbation types (**Supp. Video 1, Supp. Note 2**, & **Supp. Fig. 4**).

However, sensory brain regions like area 2 are typically characterized in terms of how neural activity encodes sensory stimuli (35,40,41). Thus, we examine whether rates inferred by AutoLFADS explain observed spikes better than a typical area 2 neural encoding model, in which neural activity is fit to some function of the state of the arm. We fit a generalized linear model (GLM) for each neuron over both active and passive movements, where the firing rate was solely a function of the position and velocity of the hand, as well as the contact forces with the manipulandum handle (35) (GLM predictions shown in **Fig. 3b**). We then compared the ability of the GLM and AutoLFADS to capture each neuron’s observed response using pseudo-R^2^ (pR^2^), a metric similar to R^2^ but adapted for the Poisson statistics of neural firing (42). For the vast majority of neurons across two datasets, AutoLFADS predicted the observed activity significantly better than GLMs (p<0.05 for 110/121 neurons, bootstrap; see *Methods*), and there were no neurons for which the GLM produced better predictions than AutoLFADS (**Fig. 3c**).

We used linear decoding to extract subspaces of neural activity that corresponded to × and y hand velocities for both smoothed spikes and rates inferred by AutoLFADS (**Fig. 3d**). The AutoLFADS rates contained subspaces that more clearly separated hand velocities for all active conditions and all passive conditions than smoothing, showing that they are better represented in the modeled dynamics of area 2. Further, single-trial hand velocity decoding from rates inferred by AutoLFADS for active trials was substantially more accurate than that of smoothing, and also more accurate than decoding from the output of any random search model (**Fig. 3e**). On a second dataset that included whole-arm motion tracking, the velocity of all joint angles was decoded from AutoLFADS rates with higher accuracy than from smoothing or GPFA (**Fig. 3f**, right; p<0.05 for all joints, paired, one-sided Student’s t-Test).

Again, these results demonstrate the dual advantages of AutoLFADS over random search in unsupervised model selection and higher peak performance across PSTH reconstruction and decoding (hand and joint velocities). In an analysis unique to this sensory area, we additionally show that AutoLFADS rates explain neural spiking better than the state of the arm.

### AutoLFADS accurately captures single-trial dynamics during cognition

We further tested the generality of AutoLFADS by applying it to data collected from dorsomedial frontal cortex (DMFC) during a cognitive time estimation task. This task requires the brain to perform both sensory input processing and internal timing. DMFC is often considered an intermediate region in the sensorimotor hierarchy (43), interfacing with both low-level sensory and motor (PMd/M1) areas. Its activity seems to relate to higher-level aspects of motor control, including motor timing (44,45), planning movement sequences (46), learning sensorimotor associations (47) and context-dependent reward modulation (48). Importantly for our experiments, population dynamics in DMFC are tied to behavioral correlates such as movement production time (15,45,49). This DMFC dataset consisted of recordings taken as a macaque attempted to reproduce a given time interval by waiting before initiating a response movement (**Fig. 4a**, left; see *Methods* for details).

**Figure 4.**
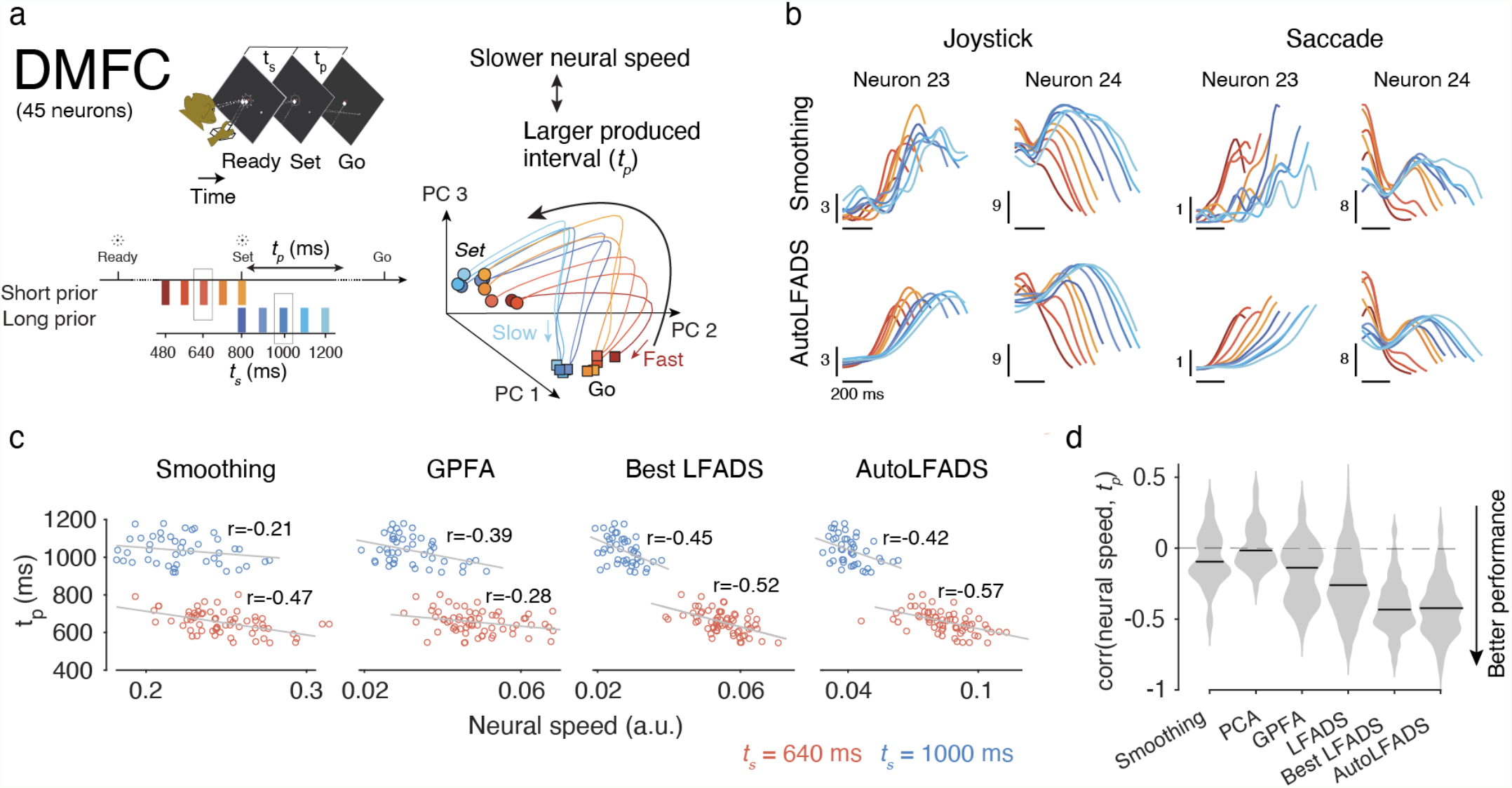
Application of AutoLFADS to data from dorsomedial frontal cortex (DMFC). (a) Top left: the time interval reproduction task. Bottom left: timing conditions used. Right: schematic illustrating the inverse correlation between neural speed and monkey’s produced time (*t*_*p*_). (b) PSTHs for <u>two</u> example neurons during the Set-Go period of rightward trials for two response modalities and all values of *t*_*s*_. <u>Vertical scale bars denote spikes / sec.</u> (c) Example plots showing correlations between neural speed and behavior (i.e., production time, *t*_*p*_) for individual trials across two timing intervals (red: 640 ms blue: 1000 ms). Neural speed was obtained based on the firing rates inferred from smoothing, GPFA, the LFADS model <u>(no CD)</u> with best median speed-*t*_*p*_ correlation across the 40 different task conditions (Best LFADS), and AutoLFADS. (d) Distributions of correlation coefficients across 40 different task conditions. Horizontal lines denote medians. For LFADS, the distribution includes correlation values for all 96 models with random HPs (40×96 values).

Consistent with our observations on M1/PMd and area 2 data, AutoLFADS-inferred rates for this dataset showed consistent, denoised structure at the single-trial level (**Supp. Fig. 2**). In particular, AutoLFADS captures the *t*_*s*_- dependent ordering of PSTHs observed in previous work for both response modalities (**Fig. 4b**) (49). We found similar results in a qualitative characterization of AutoLFADS latent factors (**Supp. Note 3 & Supp. Fig. 5**). Quantitative comparison of the PSTHs shows that AutoLFADS-inferred rates again achieved a better match to the empirical PSTHs than all of the random search models (**Supp. Fig. 2**), providing further evidence that AutoLFADS can achieve superior models without expert tuning of regularization HPs or supervised model selection criteria.

Previous studies have shown that the monkey’s produced time interval (*t*_*p*_) is negatively correlated to the speed at which the neural trajectories evolve during the Set-Go period (**Fig. 4a**, right) (15,49). To evaluate the correspondence between neural activity and behavior on a single-trial basis, we estimated neural speeds using representations produced by smoothing spikes, GPFA, principal component analysis (PCA), the best random search model (‘Best LFADS’, see *Methods* for details), and an AutoLFADS model, and measured the trial-by-trial correlation between the estimated speeds and *t*_*p*_. If a given representation of neural activity is more informative about behavior, we expect a stronger (more negative) correlation between predicted and observed *t*_*p*_.

We show correlation values for individual trials across two different timing conditions (*t*_*s*_) (**Fig. 4c**), and summarize across all 40 task conditions (**Fig. 4d**). We observed consistent negative correlations between *t*_*p*_ and the estimated neural speed from rates obtained by different methods. Correlations from rates inferred by AutoLFADS were significantly better than all unsupervised approaches (p<0.001, Wilcoxon signed rank test), and comparable with the supervised selection approach (‘Best LFADS’, p=0.758, Wilcoxon signed rank test), despite using no task information.

### Unsupervised HP tuning avoids representational and trial-average biases

Latent variable models like LFADS are inherently underdefined, so supervised model selection criteria can bias HP tuning towards models with certain types of representations (e.g., those that contain information about certain behavioral variables or trial averages), at the expense of explaining the neural activity. By comparing model performance with respect to different supervised metrics, we show that in most cases there is substantial disagreement over which models are highest-performing (**Fig. 5**). For example, selecting a random search model based on kinematic decoding for area 2 would result in suboptimal PSTH reconstruction and vice-versa. On the other hand, by selecting models using log-likelihood of observed spikes, AutoLFADS learns representations that perform well across all of our supervision metrics.

**Figure 5.**
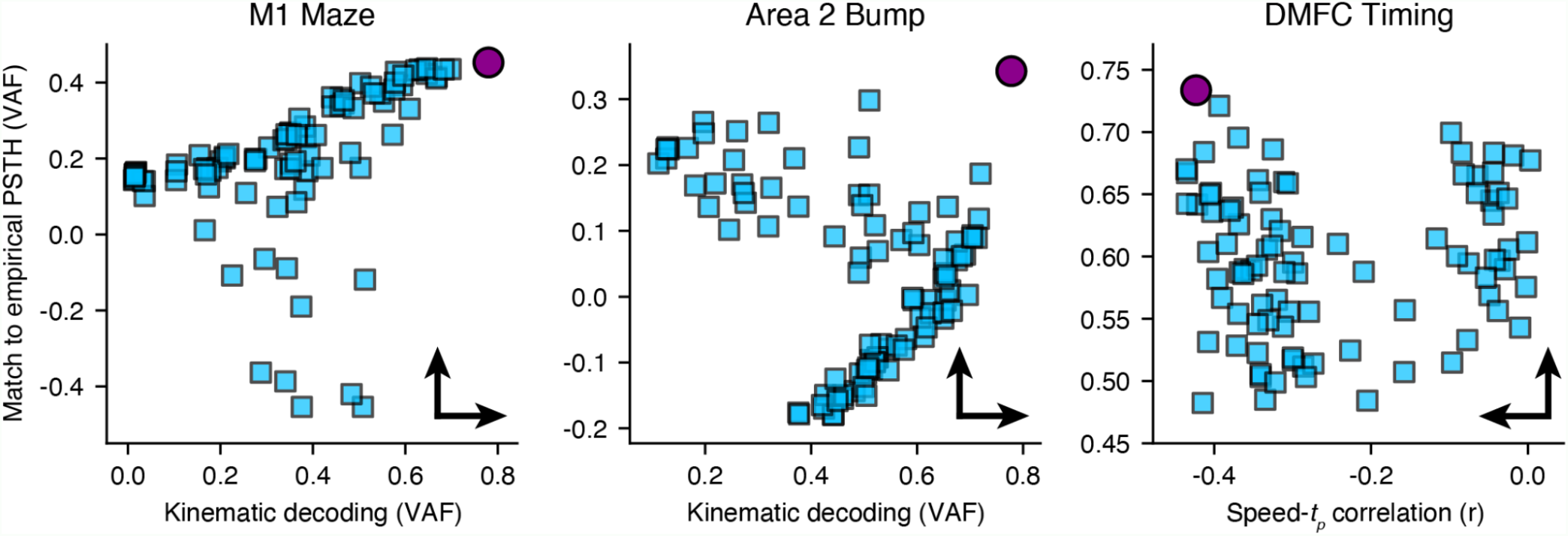
Disagreement between supervised metrics on several datasets. Panels compare the performance of AutoLFADS and a 96-model random search according to two supervised metrics for each dataset. Conventions and metric values are the same as those given in **Figs. 2d, 3e, and 4d** and **Supp. Fig. 2**). Arrows indicate the direction of better performance for each metric.

## Discussion

The original LFADS work (20) provided a method for inferring latent dynamics, denoised firing rates, and external inputs from large populations of neurons, producing representations that were more informative of behavior than previous approaches (31). However, application of LFADS to neural populations with different dynamics, strong external inputs, or unconstrained behavior would have necessitated time-consuming and subjective manual tuning. This process was made more complicated by identity overfitting, which necessitated supervised model selection metrics (e.g. decoding performance or match to empirical PSTHs). This approach to model selection is flawed for two main reasons. First, it makes model selection difficult or impossible in experiments without structured tasks or external behavioral variables. Second, it biases model selection away from accurate representation of neural activity and toward representational hypotheses or trial averages.

In the current work, we show that with a novel regularization technique and efficient hyperparameter tuning, it is possible to train high-performing LFADS models for neural spiking datasets with arbitrary size, trial structure, and dynamical complexity. By preventing overfitting to spikes, CD enables unsupervised model selection (and thereby HP tuning) for LFADS using only spiking data via validation negative log-likelihood. Before AutoLFADS, unsupervised HP tuning for LFADS was not merely difficult, but impossible. Instead, supervision was required to systematically optimize hyperparameters. This could be achieved by (1) decoding an external behavioral variable from inferred rates or (2) comparing inferred rates to empirical PSTHs. The first option required measured behavioral data, biased evaluation toward representational hypotheses, and only evaluated a handful of potentially low-variance dimensions in the population activity. The second required repeated trials, relied on hand-tuned parameters like smoothing width, and biased evaluation towards trial-averaged responses. Critically, AutoLFADS is both less restrictive and higher performing than previous approaches. It does not require repeated trials or behavioral variables for model selection, yet consistently outperforms models selected via the same supervised criteria.

We demonstrated several other properties of the AutoLFADS training approach which have broad implications. On the maze task, we showed that AutoLFADS models are more robust to dataset size, opening up new lines of inquiry on smaller datasets and reducing the number of trials that must be conducted in future experiments. Using the random target task, we demonstrated how AutoLFADS needs no task information in order to generate rich dynamical models of neural activity. This enables the study of dynamics during richer tasks and reuse of datasets collected for another purpose. With the perturbed reaching task, we demonstrated the first application of dynamical modeling, as opposed to encoder-based modeling, to the highly input-driven somatosensory area 2. Finally, in the timing task, we showed that AutoLFADS found the appropriate balance between inputs and internal dynamics for a cognitive area by modeling DMFC. It’s worth noting that in all the above scenarios, AutoLFADS models could match or outperform the best LFADS models in terms of behavioral metrics, despite not using any behavioral information during training or model selection. Some comparisons to other methods are already available in the recently introduced Neural Latents Benchmark, and more are forthcoming (29). We provide further information on running AutoLFADS in the cloud and extending PBT in **Supp. Note 4**.

AutoLFADS inherits some of the flaws of the LFADS model. For example, the linear-exponential-Poisson observation model is likely an oversimplification. However, we used this architecture as a starting point to show that a large-scale hyperparameter search is feasible and beneficial. By enabling large-scale searches, we can be reasonably confident that any performance differences achieved by future architecture changes will be due to real differences in modeling capabilities rather than a simple lack of HP optimization. AutoLFADS does still require the user to select HP ranges for initialization, but these ranges are more relevant to optimization speed than to final performance. Evolutionary processes can and do guide HP trajectories outside of the initialization ranges if this leads to higher-performing models. If the user has no prior for reasonable search ranges, we recommend using a large initialization range at the risk of requiring more compute time.

Taken together, these results show that AutoLFADS provides an accessible and extensible framework for generalized inference of single-trial neural dynamics with unprecedented accuracy that has the potential to unify the way we study computation through dynamics across brain areas and tasks.

## Data Availability

Highly similar datasets have been made publicly available in a standardized format through the Neural Latents Benchmark (29) at <u>neurallatents.github.io</u>, though recording sessions are different from those used in this paper. We prefer that readers use NLB sessions, but can make the sessions used in this paper available upon reasonable request. The random target dataset we used is publicly available at http://doi.org/10.5281/zenodo.3854034.

## Code Availability

Code and tutorials for running AutoLFADS are available at snel-repo.github.io/autolfads. An implementation of the original LFADS model used for random searches and manual tuning can be found at github.com/tensorflow/models.

## Acknowledgements

We thank K. Shenoy, M. Churchland, M. Kaufman, and S. Ryu for sharing the Monkey J Maze dataset. We also thank J. O’Doherty, M. Cardoso, J. Makin, and P. Sabes for making the random target dataset publicly available. This work was supported by the Emory Neuromodulation and Technology Innovation Center (ENTICe), NSF NCS 1835364, DARPA PA-18-02-04-INI-FP-021, NIH Eunice Kennedy Shriver NICHD K12HD073945, the Alfred P. Sloan Foundation, the Burroughs Wellcome Fund, and the Simons Foundation as part of the Simons-Emory International Consortium on Motor Control (CP), NIH NINDS R01 NS053603, R01 NS095251, and NSF NCS 1835345 (LEM), NSF Graduate Research Fellowships DGE-2039655 (ARS) and DGE-1324585 (RHC), the Center for Sensorimotor Neural Engineering and NARSAD Young Investigator grant from the Brain & Behavior Research Foundation (HS), NIH NINDS NS078127, the Sloan Foundation, the Klingenstein Foundation, the Simons Foundation, the McKnight Foundation, the Center for Sensorimotor Neural Engineering, and the McGovern Institute (MJ).

## Author Contributions

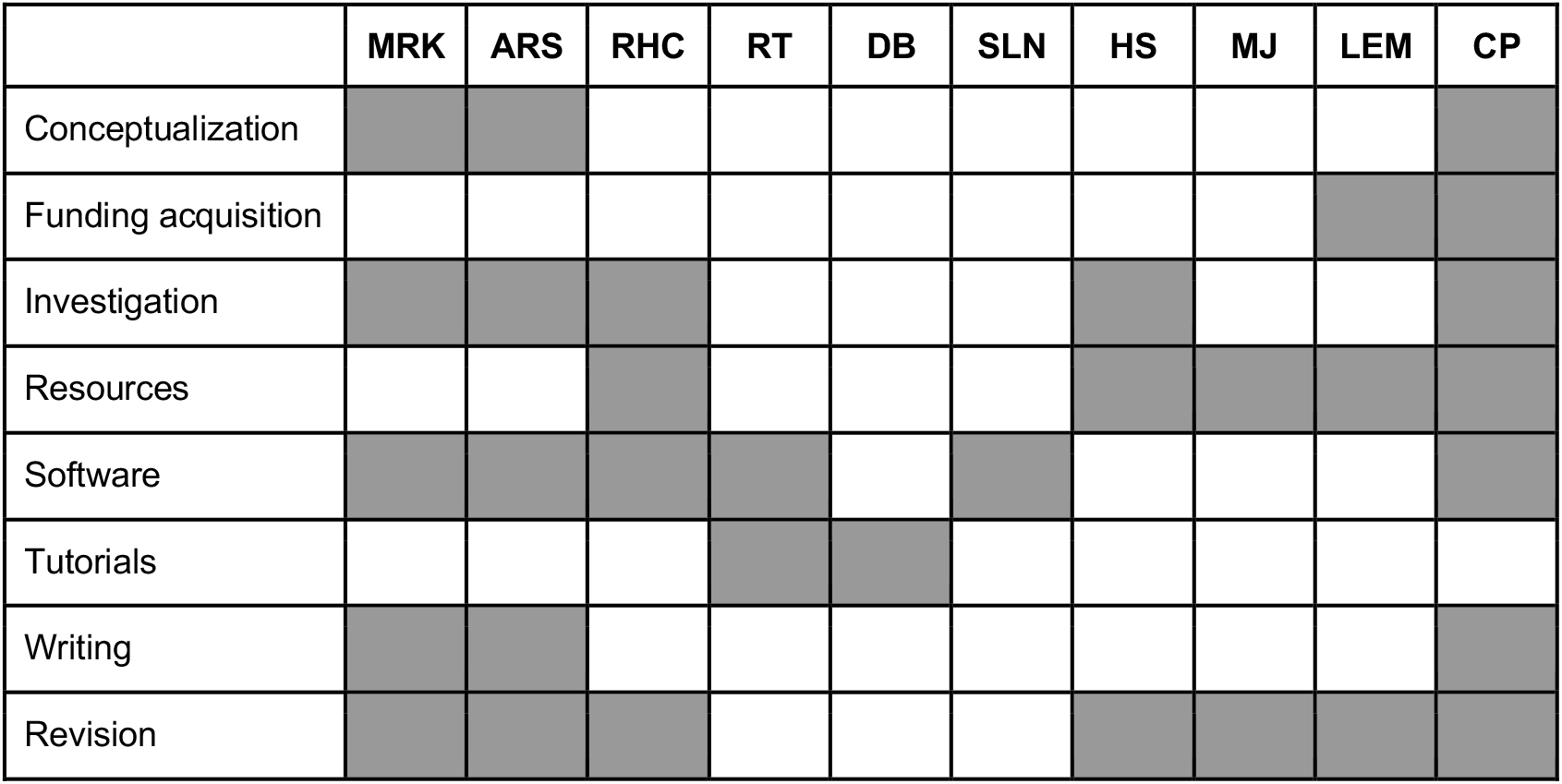

## Methods

### LFADS architecture and training

A detailed overview of the LFADS model is given in (20). Briefly: at the input to the model, a pair of bidirectional RNN encoders read over the spike sequence and produce initial conditions for the generator RNN and time-varying inputs for the controller RNN. All RNNs were implemented using gated recurrent unit (GRU) cells. The first generator state is computed with a linear mapping from the initial condition, such that the dimensionality of the inferred initial conditions is not constrained to be the same as the number of units in the generator RNN. At each time step, the generator state evolves with input from the controller and the controller receives delayed feedback from the generator. The generator states are linearly mapped to factors, which are mapped to the firing rates of the original neurons using a linear mapping followed by an exponential. The optimization objective is to minimize the negative log-likelihood of the data given the inferred firing rates, and includes KL and L2 regularization penalties.

Identical architecture and training hyperparameter values were used for most runs, with a few deviations. We used a generator dimension of 100, initial condition dimension of 100 (50 for area 2 runs), initial condition encoder dimension of 100, factor dimension of 40, controller and controller input encoder dimension of 80 (64 for DMFC runs), and controller output dimension of 4 (10 for overfitting runs).

We used the Adam optimizer with an initial learning rate of 0.01 and, for non-AutoLFADS runs, decayed the learning rate by a factor of 0.95 after every 6 consecutive epochs with no improvement to the validation loss. Training was halted for these runs when the learning rate reached 1e-5. The loss was scaled by a factor of 1e4 immediately before optimization for numerical stability. GRU cell hidden states were clipped at 5 and the global gradient norm was clipped at 200 to avoid occasional pathological training.

We used a trainable mean initialized to 0 and fixed variance of 0.1 for the Gaussian initial condition prior and set a minimum allowable variance of 1e-4 for the initial condition posterior. The controller output prior was autoregressive with a trainable autocorrelation tau and noise variance, initialized to 10 and 0.1, respectively.

Memory usage for RNNs is highly dependent on the sequence length, so batch size was varied accordingly (100 for maze and random target datasets, 500 for synthetic and area 2 datasets, and 300/400 for the DMFC dataset). KL and L2 regularization penalties were linearly ramped to their full weight during the first 80 epochs for most runs to avoid local minima induced by high initial regularization penalties. Exceptions were the runs on synthetic data, which were ramped over 70 epochs and random searches on area 2 and DMFC datasets, which used step-wise ramping over the first 400 steps.

Random searches and AutoLFADS runs used the architecture parameters described above, along with regularization HPs sampled from ranges (or initialized with constant values) given in **Supp. Table 3** and **Supp. Table 4**. Most runs used a default set of ranges, with a few exceptions outlined in the table. Dropout was sampled from a uniform distribution and KL and L2 weight HPs were sampled from log-uniform distributions. In general, more coservative hyper-HP settings (e.g., low learning rates and very wide initialization and search ranges) will work consistently across datasets. This approach is recommended in general, but as the experimenter becomes familiar with training AutoLFADS on the dataset, they may want to improve optimization speed by limiting exploration to relevant parts of the HP space.

During PBT, weights were used to control maximum and minimum perturbation magnitudes for different HPs (e.g. a weight of 0.3 results in perturbation factors between 0.7 and 1.3). The dropout and CD HPs used a weight of 0.3 and KL and L2 penalty HPs used a weight of 0.8. CD rate, dropout rate, and learning rate were limited to their specified ranges, while the KL and L2 penalties could be perturbed outside of the initial ranges. Each generation of PBT consisted of 50 training epochs. AutoLFADS training was stopped when the best smoothed validation NLL improved by less than 0.05% over the course of four generations.

Validation NLL was exponentially smoothed with α = 0.7 during training. For non-AutoLFADS runs, the model checkpoint with the lowest smoothed validation NLL was used for inference. For AutoLFADS runs, the checkpoint with the lowest smoothed validation NLL in the last epoch of any generation was used for inference. Firing rates were inferred 50 times for each model using different samples from initial condition and controller output posteriors. These estimates were then averaged, resulting in the final inferred rates for each model.

### Overfitting on synthetic data

Synthetic data were generated using a 2-input chaotic vanilla RNN (γ = 1.5) as described in the original LFADS work (20,22). The only modification was that the inputs were white Gaussian noise. In brief, the 50-unit RNN was run for 1 second (100 time steps) starting from 400 different initial conditions to generate ground-truth Poisson rates for each condition. These distributions were sampled 10 times for each condition, resulting in 4000 spiking trials. Of these trials, 80% (3200 trials) were used for LFADS training and the final 20% (800 trials) were used for validation. Detailed parameters for all datasets used in the paper can be found in **Supp. Table 2**.

We sampled 200 HP combinations from the distributions specified in **Supp. Table 3** and used them to train LFADS models on the synthetic dataset. We then trained 200 additional models with the same set of HPs using a CD rate of 0.3 (i.e., using 70% of data as input and remaining 30% for likelihood evaluation) (26). The coefficient of determination between inferred and ground truth rates was computed across all samples and neurons on the 800-sample validation set.

### M1 maze task

We used the previously-collected maze dataset (50) described in detail in the original LFADS work (20). Briefly, a male macaque monkey performed a two-dimensional center-out reaching task by guiding a cursor to a target without touching any virtual barriers while neural activity was recorded via two 92-electrode arrays implanted into M1 and dorsal PMd. The full dataset consisted of 2,296 trials, 108 reach conditions, and 202 single units.

The spiking data were binned at 1 ms and smoothed by convolution with a Gaussian kernel (30 ms s.d.). Hand velocities were computed using second order accurate central differences from hand position at 1kHz. An antialiasing filter was applied to hand velocities and all data were then resampled to 2 ms. Trials were created by aligning the data to 250 ms before and 450 ms after movement onset, as calculated in the original paper.

Datasets of varying sizes were created for LFADS by randomly selecting trials with 20, 10, and 5% of the original dataset using seven fixed seeds, and then splitting each of these into 80/20 training and validation sets for LFADS (22 total, including the full dataset). As a baseline for each data subset, we trained LFADS models with fixed HPs that had been previously found to result in high-performing models for this dataset, with the exception of controller input encoder and controller dimensionalities (see *LFADS architecture and training* and **Supp. Table 3**). We increased the dimensionality of these components to allow improved generalization to the datasets from more input-driven areas while keeping the architecture consistent across all datasets. We also trained AutoLFADS models (40 workers) on each subset using the search space given in **Supp. Table 4**. Additionally, we ran a random search using 100 HPs sampled from the AutoLFADS search space on one of the 230-trial datasets (see **Supp. Table 3**).

We used rates from spike smoothing, manually tuned LFADS models, random search LFADS models, and AutoLFADS models to predict x and y hand velocity delayed by 90 ms using ridge regression with a regularization penalty of λ = 1. Each data subset was further split into 80/20 training and validation sets for decoding. To account for the difficulty of modeling the first few time points of each trial with LFADS, we discarded data from the first 50 ms of each trial and did not use that data for model evaluation. Decoding performance was evaluated by computing the coefficient of determination for predicted and true velocity across all trials for each velocity dimension. The result was then averaged across the two velocity dimensions.

To evaluate PSTH reconstruction for random search and AutoLFADS models, we first computed the empirical PSTHs by averaging smoothed spikes from the full 2296-trial dataset across all 108 conditions. We then computed model PSTHs by averaging inferred rates across conditions for all trials in the 230-trial subset. We computed the coefficient of determination between model-inferred PSTHs and empirical PSTHs for each neuron across all conditions in the subset. We then averaged the result across all neurons.

### M1 random target task

The random target dataset consists of neural recordings and hand position data recorded from macaque M1 during a self-paced, sequential reaching task between random elements of a grid (34). For our experiments, we used only the first 30% (approx. 9 minutes) of the dataset recorded from Indy on 04/26/2016.

We started with sorted units obtained from M1 and binned their spike times at 1 ms. To avoid artifacts in which the same spikes appeared on multiple channels, we computed cross-correlations between all pairs of neurons over the first 10 sec and removed individual correlated neurons (*n* = 34) by highest firing rate until there were no pairs with correlation above 0.0625, resulting in 181 uncorrelated neurons. We remove these neurons because correlated spike artifacts can cause overfitting issues, despite the protection afforded by CD. We applied an antialiasing filter to hand velocities and smoothed the spikes by convolving with a Gaussian kernel (50 ms s.d.), a width which yields good decoding performance on this dataset. We then downsampled spikes and smoothed spikes to 2 ms to make the amount of data manageable for LFADS training. The continuous neural spiking data were chopped into overlapping segments of length 600 ms, where each segment shared its last 200 ms with the first 200 ms of the next. This overlap helps in reassembling the continuous data, as data at the ends of LFADS sequences is typically modeled better than data at the beginning. The resulting 1321 segments were split into 80/20 training and validation sets for LFADS, where the validation segments were chosen in blocks of 3 to minimize the overlap between training and validation subsets. The position data were provided at 250 Hz, so we upsampled to 500 Hz using cubic interpolation to match the neural data sampling rate.

The chopped segments were used to train an AutoLFADS model and to run a random search using 100 HPs sampled from the AutoLFADS search space (see **Supp. Tables 2 & 3**). After modeling, the chopped data were merged using a quadratic weighting of overlapping regions that placed more weight on the rates inferred at the ends of the segments. The merging technique weighted the ends of segments as *w* = 1 − *x*^2^ and the beginnings of segments as 1 − *w*, with x ranging from 0 to 1 across the overlapping points. After weights were applied, overlapping points were summed, resulting in a continuous ∼9-minute stretch of modeled data.

We computed hand velocity from position using second-order accurate central differences and introduced a 120 ms delay between neural data and kinematics. We used ridge regression (λ = 1*e* − *5*) to predict hand velocity across the continuous data using smoothed spikes, random search LFADS rates, and AutoLFADS rates. We computed coefficient of determination for each velocity dimension individually and then averaged the two velocity dimensions to compute decoding performance.

To prepare the data for subspace visualization, the continuous activity for each neuron was soft-normalized by subtracting its mean and dividing by its 90th quantile plus an offset of 0.01. Trials were identified in the continuous data as the intervals over which target positions were constant (314 trials). To identify valid trials, we computed the normalized distance from the final position. Trials were removed if the cursor exceeded 5% of this original distance or overshot by 5%. Thresholds (*n* = 100) were also created between 25 and 95% of the distance and trials were removed if they crossed any of those thresholds more than once. We then computed an alignment point at 90% of the distance from the final position for the remaining trials and labeled it as movement onset (227 trials). For each of these trials, data were aligned to 400 ms before and 500 ms after movement onset. The first principal component of AutoLFADS rates during aligned trials was computed and activation during the first 100 ms of each trial was normalized to [0,1]. Trials were rejected if activation peaked after 100 ms or the starting activation was more than 3 standard deviations from the mean. The PC1 onset alignment point was calculated as the first time that activity in the first principal component crossed 50% of its maximum in the first 100 ms (192 trials). This alignment point was used for all neural subspace analyses.

Movement-relevant subspaces were extracted by ridge regression from neural activity onto x-velocity, y-velocity, and speed. Similarly, position-relevant subspaces involved regression from neural activity onto x-position and y-position. For movement and position subspaces, neural and behavioral data were aligned to 200 ms before and 1000 ms after PC1 onset. Target subspaces were computed by regressing neural activity onto time series that represented relative target positions. As with the movement and position subspaces, the time series spanned 200 ms before to 1000 ms after PC1 onset. A boxcar window was used to confine the relative target position information to the time period spanning 0 to 200 ms after PC1 onset, and the rest of the window was zero-filled. For kinematic prediction from neural subspaces, we used a delay of 120 ms and 80/20 trial-wise training and validation split. For each behavioral variable and neural data type, a 5-fold cross-validated grid search (*n* = 100) was used on training data to find the best-performing regularization across orders of magnitude between 1e-5 and 1e4.

Single subspace dimensions were aligned to 200 ms before and 850 ms after PC1 onset for plotting. Subspace activations were calculated by computing the norm of activations across all dimensions of the subspace and then rescaling the min and max activations to 0 and 1, respectively. Multidimensional subspace plots for the movement subspace were aligned to 180 ms before and 620 ms after PC1 onset and for target subspace 180 ms before and 20 ms after.

### Area 2 bump task

The sensory dataset consisted of two recording sessions during which a monkey moved a manipulandum to direct a cursor towards one of eight targets (active trials). During passive trials, the manipulandum induced a mechanical perturbation to the monkey’s hand prior to the reach. Activity was recorded via an intracortical electrode array embedded in Brodmann’s area 2 of the somatosensory cortex. For the second session, joint angles were calculated from motion tracking data collected throughout the session. The first session was used for PSTH, GLM, subspace, and velocity decoding analyses and the second session was only used for pseudo-R^2^ comparison to GLM and joint angle decoding. More details on the task and dataset are given in the original paper (35).

For both sessions, only sorted units were used. Spikes were binned at 1 ms and neurons that were correlated over the first 1000 sec were removed (*n* = 2 for each session) as described for the random target task, resulting in 53 and 68 neurons in the first and second sessions, respectively. Spikes were then rebinned to 5 ms and the continuous data were chopped into 500 ms segments with 200 ms of overlap. Segments that did not include data from rewarded trials were discarded (kept 9,626 for the first session and 7,038 for the second session). A subset of the segments (30%) were further split into training and validation data (80/20) for LFADS. An AutoLFADS model (32 workers) was trained on each session and a random search (96 models) was performed on the first session (see **Supp. Table 3 and Supp. Table 4**). After modeling, LFADS rates were then reassembled into their continuous form, with linear merging of overlapping data points.

Empirical PSTHs were computed by convolving spikes binned at 1 ms with a half-Gaussian (10 ms s.d.), rebinning to 5 ms, and then averaging across all trials within a condition. LFADS PSTHs were computed by similarly averaging LFADS rates. Passive trials were aligned 100 ms before and 500 ms after the time of perturbation, and active trials were aligned to the same window around an acceleration-based movement onset (35). Neurons with firing rates lower than 1 Hz were excluded from the PSTH analysis. To quantitatively evaluate PSTH reconstruction, the coefficient of determination was computed for each neuron and passive condition in the four cardinal directions, and these numbers were averaged for each model.

As a baseline for how well AutoLFADS could reconstruct neural activity, we fit generalized linear models (GLMs) to each individual neuron’s firing rate, based on the position and velocity of and forces on the hand (see (35) for details of the hand kinematic-force GLM). Notably, in addition to fitting GLMs using the concurrent behavioral covariates, we also added 10 bins of behavioral history (50 ms) to the GLM covariates, increasing the number of GLM parameters almost tenfold. Furthermore, because we wanted to find the performance ceiling of a behavioral-encoder-based GLMs to compare with the dynamics-based AutoLFADS, we purposefully did not cross-validate the GLMs. Instead, we simply evaluated GLM fits on data used to train the model.

To evaluate AutoLFADS and GLMs individually, we used the pseudo-R^2^ (pR^2^), a goodness-of-fit metric adapted for the Poisson-like statistics of neural activity. Like variance-accounted-for and R^2^, pR^2^ has a maximum value of 1 when a model perfectly predicts the data, and a value of 0 when a model predicts as well as a single parameter mean model. Negative values indicate predictions that are worse than a mean model. For each neuron, we compared the pR^2^ of the AutoLFADS model to that of the GLM (**Fig. 5e**). To determine statistically whether AutoLFADS performed better than GLMs, we used the relative-pR^2^ (rpR^2^) metric, which compares the two models against each other, rather than to a mean model (see (51) for full description of pR^2^ and rpR^2^). In this case, a rpR^2^ value above 0 indicated that AutoLFADS outperformed the GLM (indicated by filled circles in **Fig. 5e**). We assessed significance using a bootstrapping procedure, after fitting both AutoLFADS and GLMs on the data. On each bootstrap iteration, we drew a number of trials from the session (with replacement) equal to the total number of trials in the session, evaluating the rpR^2^ on this set of trials as one bootstrap sample. We repeated this procedure 100 times. We defined neurons for which at least 95 of these rpR^2^ samples were greater than 0 as neurons that were predicted better by AutoLFADS than a GLM. Likewise, neurons for which at least 95 of these samples were below 0 would have been defined as neurons predicted better by GLM (though there were no neurons with this result).

For the subspace analysis, spikes were smoothed by convolution with a Gaussian (50 ms s.d.) and then rebinned to 50 ms. Neural activity was scaled using the same soft-normalization approach outlined for the random target task subspace analysis. Movement onset was calculated using the acceleration-based movement onset approach for both active and passive trials. For decoder training, trials were aligned to 100 ms before to 600 ms after movement onset. For plotting, trials were aligned to 50 ms before and 600 ms after movement onset. The data for successful reaches in the four cardinal directions was divided into 80/20 trial-wise training and validation partitions. Separate ridge regression models were trained to predict each hand velocity dimension for active and passive trials using neural activity delayed by 50 ms (total 4 decoders). The regularization penalty was determined through a 5-fold cross validated grid search of 25 values from the same range as the random target task subspace decoders.

For hand velocity decoding, spikes during active trials were smoothed by convolution with a half-Gaussian (50 ms s.d.) and neural activity was delayed by 100 ms relative to kinematics. The data were aligned to 200 ms before and 1200 ms after movement onset and trials were split into 80/20 training and validation sets. Simple regression was used to estimate kinematics from neural activity and the coefficient of determination was computed and averaged across x- and y-velocity.

GPFA was performed on segments from all rewarded trials using a latent dimension of 20 and Gaussian smoothing kernel (30 ms s.d.). Decoding data were extracted by aligning data from active trials to 200 ms before and 500 ms after movement onset. Data were split into 80/20 training and validation sets and neural activity was lagged 100 ms behind kinematics. Ridge regression (λ = 0.001) was used to decode all joint angle velocities from smoothed spikes (half-Gaussian, 50 ms kernel s.d.), rates inferred by GPFA, and rates inferred by AutoLFADS.

### DMFC timing task

The cognitive dataset consisted of one session of recordings from the dorsomedial frontal cortex (DMFC) while a monkey performed a time interval reproduction task. The monkey was presented with a “Ready” visual stimulus to indicate the start of the interval and a second “Set” visual stimulus to indicate the end of the sample timing interval, *t*_*s*_. Following the Set stimulus, the monkey made a response (“Go”) so that the production interval (*t*_*p*_) between Set and Go matches the corresponding *t*_*s*_. The animal responded with either a saccadic eye movement or a joystick manipulation to the left or right depending on the location of a peripheral target. The two response modalities, combined with 10 timing conditions (*t*_*s*_) and two target locations, led to a total of 40 task conditions. A more detailed description of the task is available in the original paper (49).

To prepare the data for LFADS, the spikes from sorted units were binned at 20 ms. To avoid artifacts from correlated spiking activity, we computed cross-correlations between all pairs of neurons for the duration of the experiment and sequentially removed individual neurons (*n* = 8) by the number of above-threshold correlations until there were no pairs with correlation above 0.2, resulting in 45 uncorrelated neurons. Data between the “Ready” cue and the trial end was chopped into 2600 ms segments with no overlap. The first chop for each trial was randomly offset by between 0 and 100 ms to break any link between trial start times and chop start times. The resulting neural data segments (1659 total) were split into 80/20 training and validation sets for LFADS. An AutoLFADS model (32 workers) and random search (96 models) were trained on these segments (see **Supp. Table 3** and **Supp. Table 4**).

For all analyses of smoothed spikes, smoothing was performed by convolving with a Gaussian kernel (widths described below) at 1 ms resolution.

Empirical PSTHs were computed by trial-averaging smoothed spikes (25 ms kernel s.d., 20 ms bins) within each of the 40 conditions. LFADS PSTHs were computed by similarly averaging LFADS rates. The coefficient of determination was computed between inferred and empirical PSTHs across all neurons and time steps during the “Ready-Set” and “Set-Go” periods for each condition and then averaged across periods and conditions.

To visualize low-dimensional neural trajectories, demixed principal component analysis (dPCA; (12)) was performed on smoothed spikes (40 ms kernel s.d., 20 ms bins) and AutoLFADS rates during the “Ready-Set” period. The two conditions used were rightward and leftward hand movements with *t* _*s*_ = 1000 *ms*.

Besides LFADS/AutoLFADS, three alternate methods were applied for speed-tp correlation comparisons: spike smoothing, GPFA, and PCA. For spike smoothing, analyses were performed by smoothing with a 40 ms s.d.. For GPFA, a model was trained on the concatenated training and validation sets with a latent dimension of 9. Principal component analysis (PCA) was performed on smoothed spikes (40 ms kernel s.d., 20 ms bins) and 5-7 top PCs that explained more than 75% of data variance across conditions were included in the later analysis.

Neural speed was calculated by computing distances between consecutive time bins in a multidimensional state space and then averaging the distances across the time bins for the production epoch. The number of dimensions used to compute the neural speed was 45, 5-7, 9, and 45 for smoothing, PCA, GPFA and LFADS, respectively. The Pearson’s correlation coefficient between neural speed and the produced time interval was computed across trials within each condition.

