## Supplementary Material for "A large-scale neural network training framework for generalized estimation of single-trial population dynamics"

| Dataset | Subset | Metric | Smoothing | GPFA | Poisson GLM | LFADS (Agg.) | LFADS (Best) | LFADS-CD (Agg.) | LFADS-CD (Best) | AutoLFADS |
|---|---|---|---|---|---|---|---|---|---|---|
| Synthetic | | Rate recon. ($R^2$) | | | | 0.635 | 0.808 | 0.728 | **0.827** | |
| M1 Maze | 92-trial (manual) | HVD ($R^2$) | 0.611* | | | | 0.260* | | | **0.673*** |
| | 184-trial (manual) | HVD ($R^2$) | 0.637* | | | | 0.482* | | | **0.755*** |
| | 368-trial (manual) | HVD ($R^2$) | 0.640* | | | | 0.652* | | | **0.839*** |
| | 1836-trial (manual) | HVD ($R^2$) | 0.675* | | | | 0.880* | | | **0.901*** |
| | 184-trial (search) | HVD ($R^2$) | 0.605 | 0.627 | | 0.352 | 0.698 | | | **0.780** |
| | | PSTH recon. ($R^2$) | | | | 0.192 | 0.437 | | | **0.453** |
| M1 RTT | | HVD ($R^2$) | 0.524 | 0.509 | | 0.530 | 0.751 | | | **0.757** |
| Area 2 Bump | Session 1 | HVD ($R^2$) | 0.517 | 0.555 | | 0.510 | 0.720 | | | **0.778** |
| | | PSTH recon. ($R^2$) | | | | 0.034 | 0.298 | | | **0.343** |
| | | Spike pred. ($pR^2$) | | | 0.005** | | | | | **0.073**** |
| | Session 2 | JVD-SA ($R^2$) | 0.189* | 0.214* | | | | | | **0.284*** |
| | | JVD-SR ($R^2$) | 0.364* | 0.388* | | | | | | **0.477*** |
| | | JVD-SF ($R^2$) | 0.506* | 0.521* | | | | | | **0.668*** |
| | | JVD-EF ($R^2$) | 0.446* | 0.444* | | | | | | **0.613*** |
| | | JVD-RP ($R^2$) | 0.041* | 0.057* | | | | | | **0.084*** |
| | | JVD-WF ($R^2$) | 0.091* | 0.102* | | | | | | **0.243*** |
| | | JVD-WA ($R^2$) | 0.027* | 0.030* | | | | | | **0.089*** |
| | | Spike pred. ($pR^2$) | | | 0.005** | | | | | **0.096**** |
| DMFC Timing | | Speed-tp corr. (r) | -0.103** | -0.135** | | -0.259** | **-0.435**** | | | -0.423** |
| | | PSTH recon. ($R^2$) | | | | 0.591 | 0.721 | | | **0.734** |

**Supplementary Table 1.** Collected performance metrics across all datasets. Metrics include coefficient of determination (R2), pseudo-R2 (pR2), and correlation coefficient (r). Hand velocity decoding is denoted by HVD and joint velocity decoding by JVD. Joint angle abbreviations are as described in Fig. 3f. Asterisks denote aggregated metrics from multiple models, with * indicating mean and ** indicating median.

| Dataset | Bin size (ms) | Window (bins) | Overlap (bins) | Units | Train samples | Valid samples | Total samples |
|---|---|---|---|---|---|---|---|
| Synthetic | 10 | 100 | 0 | 50 | 3200 | 800 | 4000 |
| M1 Maze | 2 | 350 | 0 | 202 | 92, 184, 368, 1836 | 23, 46, 92, 460 | 2396 |
| M1 RTT | 2 | 300 | 100 | 181 | 1057 | 264 | 1321 |
| Area 2 (sess. 1) | 5 | 100 | 40 | 53 | 2310 | 578 | 9626 |
| Area 2 (sess. 2) | 5 | 100 | 40 | 68 | 1581 | 396 | 7038 |
| DMFC | 20 | 130 | 0 | 45 | 1327 | 332 | 1659 |

**Supplementary Table 2.** Dataset parameters.

| | NLL | NUM | LR | CD | DO | KL CO | KL IC | L2 Con | L2 Gen | L2 IC Enc | L2 CI Enc |
|---|---|---|---|---|---|---|---|---|---|---|---|
| Defaults | MEAN | 100 | 1e-2 | 0 | (0.0, 0.6) | (1e-6, 1e-4) | (1e-6, 1e-3) | (1e-4, 1e0) | (1e-4, 1e0) | 0 | 0 |
| Overfitting | | 200 | | 0 or 0.3 | (0.0, 0.7) | (1e-6, 1e-3) | | (1e-5, 1e0) | (1e-5, 1e0) | (1e-5, 1e-3) | (1e-5, 1e-3) |
| Maze | | | | | | | | | | | |
| Maze (manual) | SUM | 1 | | 0 | 0.02 | 1 | 1 | 10 | 10 | | |
| RTT | | | | | | | | | | | |
| Area2 | SUM | 96 | | | | (1e-1, 1e1) | (1e-1, 5e0) | (1e0, 3e4) | (1e0, 3e4) | | |
| DMFC | SUM | 96 | (1e-5, 7e-3) | | | (1e-1, 1e1) | (1e-1, 5e0) | (1e0, 3e4) | (1e0, 3e4) | | |

**Supplementary Table 3.** Hyperparameter ranges for LFADS runs and random searches. Blank cells or missing rows indicate use of default values. NLL indicates whether the NLL was aggregated by mean or sum across units and time points. NUM is the number of models trained. LR is the learning rate. CD is the coordinated dropout rate (i.e. proportion of samples dropped at input). DO is the dropout rate. KL indicates weight applied to KL divergence of a posterior from its prior. CO indicates controller output distributions and IC indicates initial condition distributions. L2 indicates weight applied to the Frobenius norm of the recurrent kernel of the GRU cell. Con indicates the controller GRU cell, Gen indicates the generator GRU cell, IC Enc indicates the initial condition encoder GRU cells, and CI Enc indicates the controller input encoder GRU cells.

| | NLL | NUM | | LR | CD | DO | KL CO | KL IC | L2 Con | L2 Gen |
|---|---|---|---|---|---|---|---|---|---|---|
| Defaults | MEAN | 40 | Ranges | (1e-5, 1e-2) | (0.01, 0.7) | (0.0, 0.6) | (1e-6, 1e-4) | (1e-6, 1e-3) | (1e-4, 1e0) | (1e-4, 1e0) |
| | | | Initial values | 1e-2 | 0.5 | | | | | |
| | | | Explore weight | 0.3 | 0.3 | 0.3 | 0.8 | 0.8 | 0.8 | 0.8 |
| | | | Sampling | - | - | uniform | loguniform | loguniform | loguniform | loguniform |
| | | | Limit explore | TRUE | TRUE | TRUE | FALSE | FALSE | FALSE | FALSE |
| Maze | | | Ranges | | | | | | | |
| | | | Initial values | | | | | | | |
| RTT | | | Ranges | | | | | | | |
| | | | Initial values | | | | | | | |
| Area2 | | 32 | Ranges | (1e-5, 5e-3) | | | | (1e-6, 1e-4) | | |
| | | | Initial values | 4e-3 | | | | | | |
| DMFC | | 32 | Ranges | (1e-5, 7e-3) | (0.01, 0.99) | | | (1e-5, 1e-3) | | |
| | | | Initial values | 5e-3 | | | | | | |

**Supplementary Table 4.** Hyperparameter ranges for AutoLFADS runs. Conventions are the same as in **Supp. Table 2**.

**Supplementary Video 1.** Projection of four-dimensional inputs, from -100 ms to 200 ms around movement onset, into the top three principal components across all conditions, with separate plots for each movement direction. Darker lines indicate active trials while lighter lines denote passive trials. Large dots indicate average initial input in PC space. Thick line indicates average input trace for each direction, indicated by color, while thin colored lines show input traces for ten randomly chosen trials. Vertical scale bar is a.u.

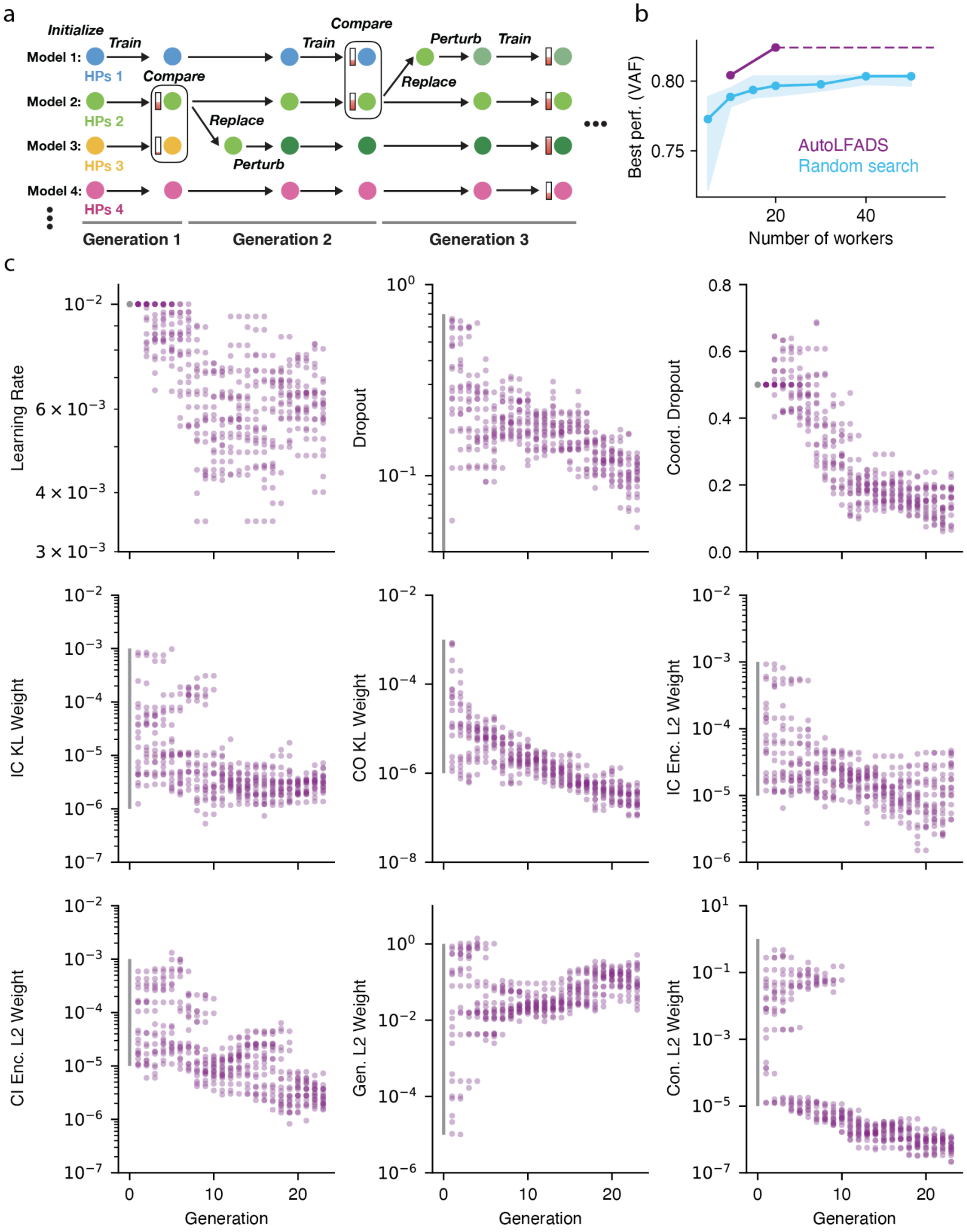


**Supplementary Figure 1 | Training AutoLFADS models with Population-Based Training.** (a) Schematic of the PBT approach to HP optimization. Each colored circle represents an LFADS model with a certain HP configuration and partially filled bars represent model performance (higher is better). In our case, performance is measured by exponentially-smoothed validation log-likelihood at the end of each generation. Models are trained for fixed intervals (generations), between which poorly-performing models are replaced by copies of better-performing models with perturbed HPs. (b) True rate recovery performance of AutoLFADS vs. best random search model (no CD) for a given number of workers. Random searches were simulated by drawing from the pool of runs shown in **Fig. 1c**. Shaded regions denote upper and lower quartiles for 100 draws. (c) Hyperparameter progressions for the 20-worker AutoLFADS run shown in the previous panel. Initialization values are shown as gray points and initialization ranges are shown as gray lines.

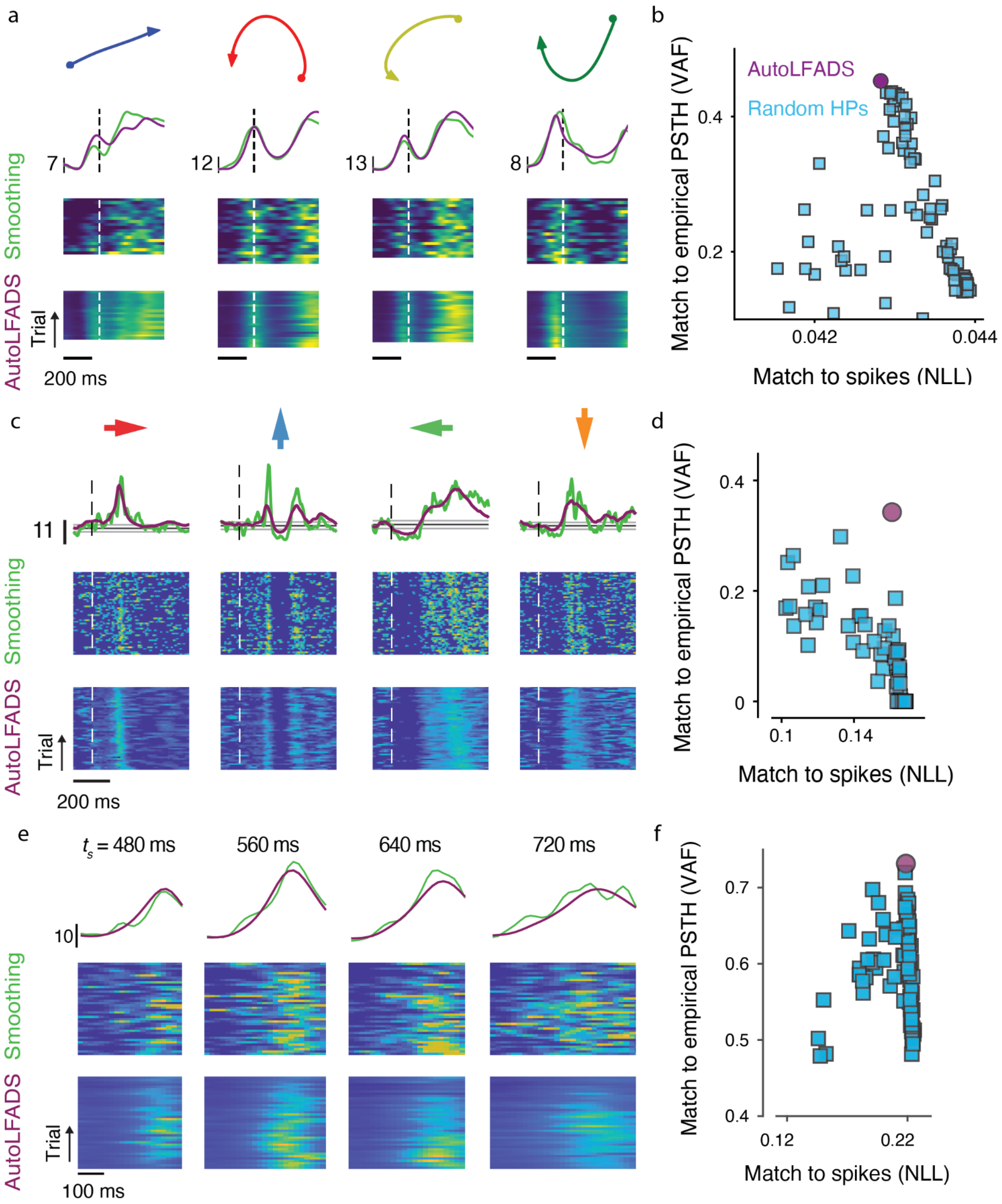


**Supplementary Figure 2 | Single-trial and PSTH recovery in diverse brain areas.** Results are shown for M1 (a, b), Area 2 (c, d) and DMFC (e, f). (a) Average reach trajectories (top), PSTHs (second row) and single-trial firing rates (bottom) obtained by smoothing (Gaussian kernel, 30 ms s.d.) or AutoLFADS for a single neuron across 4 reach conditions. Data is modeled at 2 ms bins. Dashed lines indicate movement onset and vertical scale bars denote rates (spikes/s). (b) Performance in replicating the empirical PSTHs computed on all trials using rates inferred from a 184-trial training set using AutoLFADS and LFADS with random HPs (100 models; no CD). (c) PSTHs and single-trial firing rates for a single neuron across 4 passive perturbation directions. Smoothing was performed using a Gaussian kernel with 10 ms s.d.. Dashed lines indicate movement onset. (d) Comparison of AutoLFADS vs. random search (no CD) in matching empirical PSTHs. (e) PSTHs and single-trial firing rates for an example neuron during the Set-Go period of leftward saccade trials across 4 different values of $t_s$ (vertical scale bar: spikes/sec). Smoothing was performed using a Gaussian kernel with 25 ms s.d.. (f) Performance in replicating the empirical PSTHs.

### Supplementary Note 1

#### AutoLFADS uncovers motor cortical dynamics without structured trials

In support of the hypothesis that AutoLFADS is picking up on meaningful dynamics that occurred throughout the session, we found that the firing rates inferred by AutoLFADS were informative of the previously-hypothesized computational role of motor cortical dynamics - i.e., linking the process of movement preparation and execution - despite the model being trained without information about the monkey's behavior (**Supp. Fig. 3**). In particular, firing rates contained subspaces that were highly informative about hand position, hand velocity, and reach target on individual trials (**Supp. Fig. 3a**) and showed clear structure relative to the task (**Supp. Fig. 3b**). To find the subspaces, we used linear regression to project neural activity onto variables related to movement goals (reach target) and movement details (position, velocity and speed). Notably, the subspace reflecting reach target was transiently active around the time of movement execution, consistent with previous studies that have demonstrated the presence of preparatory activity in motor cortex, yet revealed without an explicit preparatory period. It is likely that the rates inferred by AutoLFADS also contain yet undiscovered subspaces and representations that can be explored in this same dataset without experiments explicitly designed to reveal them. Thus, AutoLFADS has the potential to greatly improve the utility and versatility of rich behavioral datasets via a unique unsupervised modeling process (1–3).

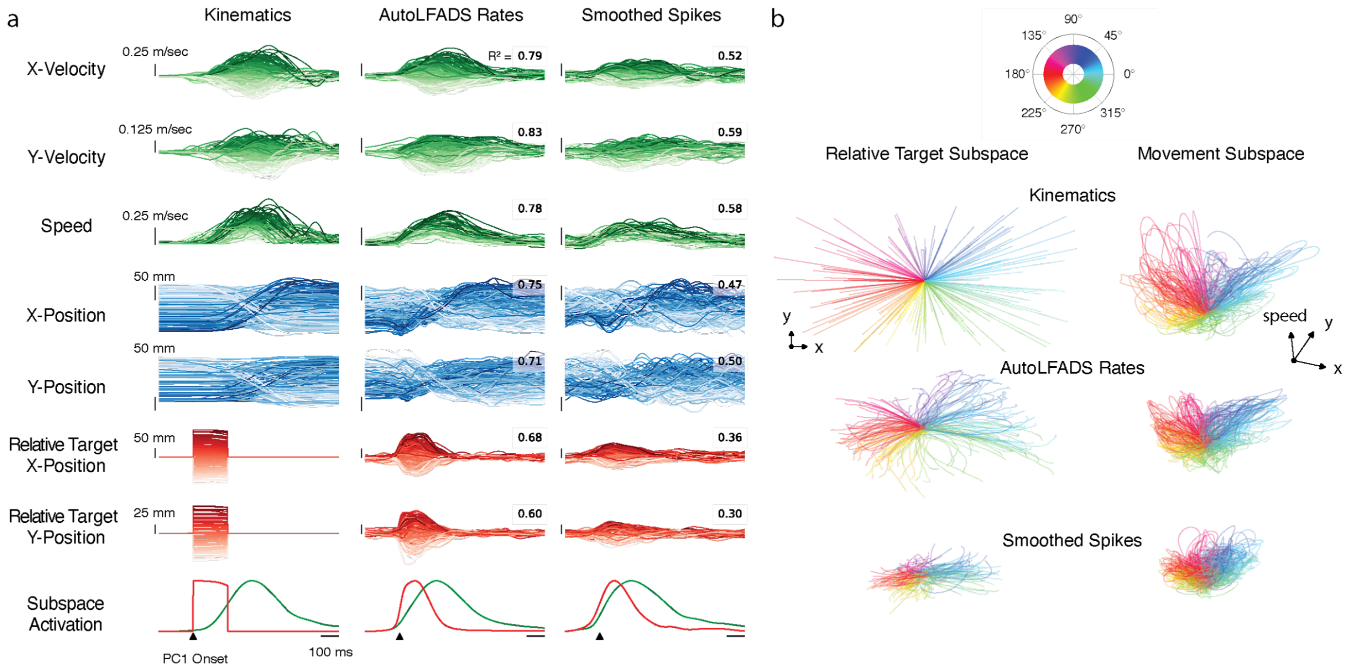


**Supplementary Figure 3 | Inferred firing rates contain neural subspaces that are informative about movement kinematics and reach targets.** (a) Kinematic and relative target variables and their corresponding neural representations, uncovered via linear regression. The quality of each projection is quantified by accuracy in decoding kinematic and target variables ($R^2$). Plots are colored by x and y distance to target, except for speed which is colored by peak speed. Bottom row represents the normalized activation of movement (green) and relative target (red) subspaces, illustrating the more transient activation in the target subspace. (b) Movement and relative target subspaces plotted as 3D trajectories and colored by angle to target.

### Supplementary Note 2

#### AutoLFADS models input-driven dynamical systems with meaningful inference of inputs

Since area 2 plays a significant role in processing sensory inputs, it stands to reason that the inputs inferred by AutoLFADS are important for successfully modeling the area's activity as a dynamical system. If AutoLFADS is successfully modeling area 2 as an input-driven dynamical system, we should expect the inferred inputs to be consistent across trials with the same behavioral conditions. In these experiments, AutoLFADS models the data as fixed-length segments without regard to trial boundaries, so there is no guarantee of the consistency of the meaning of a given input between different trials of the same condition or even within a single trial.

Despite the unsupervised modeling process, AutoLFADS inferred input trajectories that were consistent with the supervised notions of trials, directions, and perturbation types (**Supp. Fig. 4a**). Inputs were continuous over the course of a trial, implying that the model was able to pick up on statistical similarities between adjacent segments. The model also produced similar input patterns within a given condition, showing that it was able to detect the statistical patterns of a given condition from arbitrary segments of time during arbitrary trials. Finally, AutoLFADS produced distinct and logically consistent output patterns for active and passive trials. Inputs for abrupt passive movements generally had a much shorter time course that unfolded post-perturbation, while inputs for active trials began before movement and evolved more slowly (**Supp. Fig. 4a**). Visualization of these inputs highlights AutoLFADS's ability to infer distinct inputs for distinct subsets of the data (**Supp. Fig. 4b**). We highlight the time-course and 3D structure of the inferred inputs in **Supp. Video 1**.

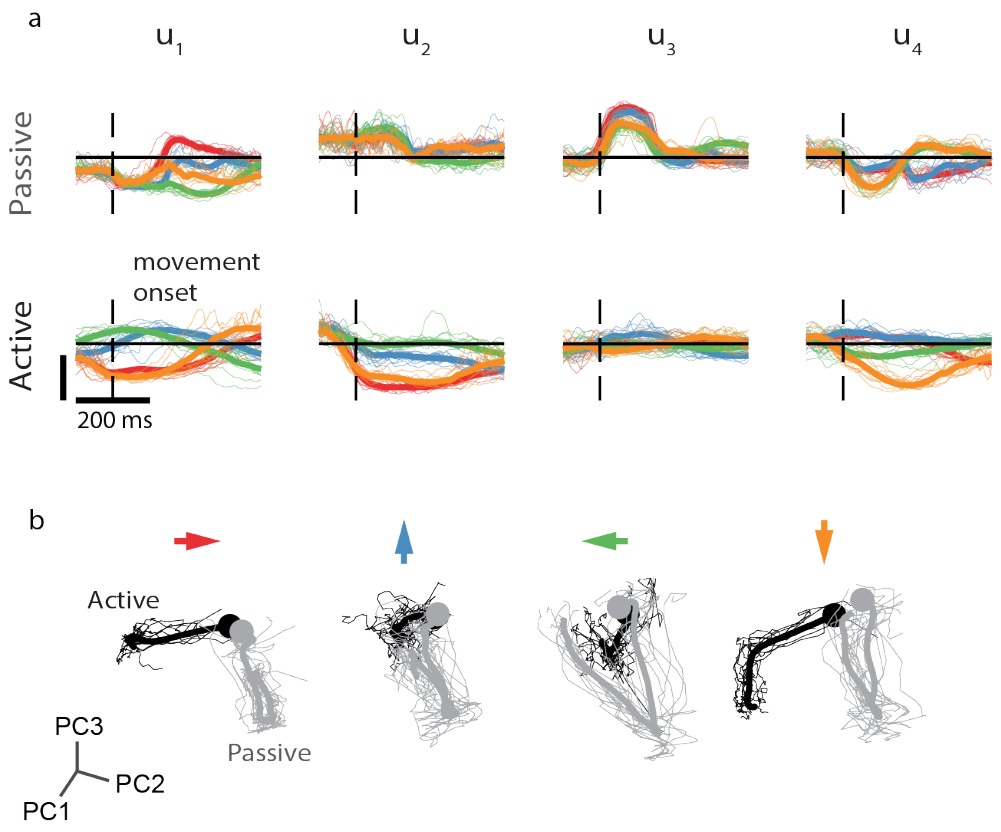


**Supplementary Figure 4 | AutoLFADS-inferred inputs for area 2 neural activity.** (a) Time-courses of the four inferred generator input dimensions for passive (top) and active (bottom) conditions. Thick line indicates average input trace for each direction, indicated by color, while thin colored lines show input traces for ten randomly chosen trials. Vertical scale bar is A.U. (b) Projection of four-dimensional inputs, from -100 ms to 200 ms around movement onset, into the top three principal components across all conditions, with separate plots for each movement direction. Darker lines indicate active trials while lighter lines denote passive trials. Large dots indicate average initial input in PC space. Thick and thin lines follow conventions in (a).

### Supplementary Note 3

#### AutoLFADS infers latent states consistent with previous findings on time interval reproduction in DMFC

Most of our evaluation on the DMFC dataset has relied on the finding that neural speed during the Set-Go period is negatively correlated with the produced time interval (4). To further explore the representation of DMFC activity learned by AutoLFADS, we visualize the top-3 PCs of the 40-D, trial-averaged latent factors. We show these PCs for all 10 timing conditions for both response modalities during both Ready-Set and Set-Go periods (**Supp. Fig. 5**). We find that during the Ready-Set period, latent states follow largely similar trajectories for all conditions, but move faster for short prior conditions. Trajectories are largely identical within each prior, which reflects the fact that the monkey doesn't know the timing condition during this period. During the Set-Go period, latent states follow similar paths traversed at different speeds for different timing conditions. All of these findings are consistent across both response modalities and reproduce previous work (4).

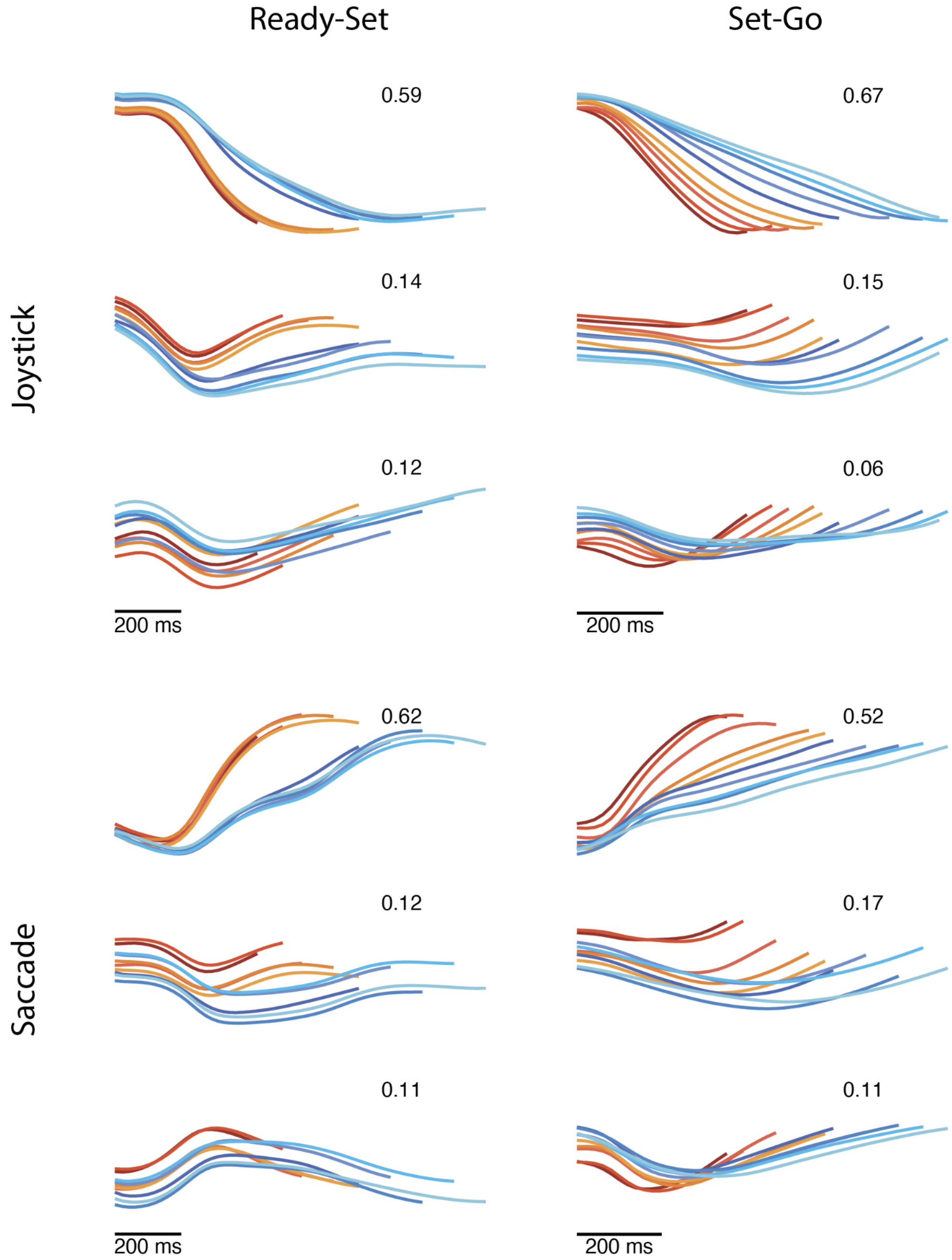


**Supplementary Figure 5 | AutoLFADS-inferred latent states for DMFC.** The top-3 PCs of trial-averaged latent states during the Ready-Set and Set-Go periods for rightward movements using both response modalities. Traces are colored by timing condition, as indicated in Fig. 4a. Plots are annotated with the fraction of variance explained by each PC.

### Supplementary Note 4

#### Running AutoLFADS in the Cloud

A key challenge with emerging, computationally-intensive data analysis methods is that the computational infrastructure and expertise necessary to make effective use of these tools is a significant barrier to widespread adoption (5). For example, many labs do not have the resources necessary to train dozens of models in parallel across many GPUs. To address this hurdle, we provide an open-source implementation of AutoLFADS designed to operate on Google Cloud Platform (GCP). Additionally, we provide a comprehensive tutorial to help novice users get started running AutoLFADS on GCP without expert knowledge of cloud computing or machine learning. The tutorial describes how to set up the framework, prepare input data, set up AutoLFADS runs, and load the final results. Users of AutoLFADS on GCP don't need to worry about the upfront hardware and labor costs associated with maintaining a local computing cluster, yet have access to virtually unlimited computation on demand. This framework allows researchers to spend less time doing non-research tasks like dependency management and hyperparameter optimization, while giving them confidence that their models are performing well, regardless of brain area or task. Our GCP implementation allows users to apply AutoLFADS without needing to purchase and maintain a local cluster. We estimate that the compute cost for a typical AutoLFADS run (including HP optimization) on GCP is between $5-25, depending on dataset and model sizes. We have created detailed tutorials to guide novice users through the setup, model training, and data retrieval processes, making AutoLFADS accessible to anyone who works with neural spiking data. We include links to the code and tutorial in *Code Availability*.

#### Extending Population-based Training

AutoLFADS performed well using simple binary tournament exploitation and perturbation exploration strategies for PBT (6). Future work might investigate alternate exploitation or exploration strategies, or whether more powerful and efficient PBT variants (7) can increase speed and performance of AutoLFADS while lowering computational cost. A current limitation of AutoLFADS is its inability to explore hyperparameters that modify the underlying model architecture. Thus, another avenue for further work lies in combining AutoLFADS with the recent techniques for automated neural architecture search (8).
